# GABA co-release guides the functional maturation of glycinergic synapses in an auditory sound localization circuit

**DOI:** 10.64898/2025.12.01.691682

**Authors:** Jongwon Lee, Brian Brockway, Karl Kandler

**Author notes:** Karl Kandler **Email:**. **Author Contributions:** J.L., B.B., and K.K. designed research; J.L. and B.B. performed research; J.L analyzed data, J.L wrote the first draft of the paper, KK wrote the final version of the manuscript. **Competing Interest Statement:** The authors have no competing interests.

## Abstract

In the mammalian brainstem and spinal cord, glycine is the primary inhibitory neurotransmitter. However, during development, many glycinergic neurons also co-release the inhibitory neurotransmitter gamma-aminobutyric acid (GABA). Although the acute effects of GABA co-release on immature synaptic transmission have been increasingly characterized, its role in synapse maturation and circuit formation remains poorly understood. Here, we investigated the developmental roles of GABA co-release at glycinergic synapses from the medial nucleus of the trapezoid body (MNTB) to the lateral superior olive (LSO), an auditory pathway essential for binaural integration and sound localization. During the first two postnatal weeks, MNTB-LSO synapses co-release GABA and undergo pronounced synaptic and circuit refinement. Using conditional knockout mice with severely diminished GABA co-release from MNTB neurons, we found that key aspects of circuit refinement, including synaptic silencing and strengthening, occurred normally. However, a disruption of GABA co-release resulted in significantly larger quantal amplitudes and a reduced readily releasable vesicle pool, impairing the high fidelity and temporal precision of synaptic transmission, which are essential for accurate binaural processing. These results reveal a critical developmental role for GABA co-release in shaping the functional synaptic architecture of glycinergic synapses involved in sound localization.

**Significance Statement:** Glycinergic neurons in the brainstem and spinal cord often co-release GABA during development, but the role of this co-release in circuit refinement or synapse maturation remains poorly understood. This study found that disruption of developmental GABA co-release at the MNTB-LSO synapse, a key part of the sound localization circuit, did not affect topographic refinement by synapse elimination or strengthening. However, impaired GABA co-release prevented synapses from developing the characteristic features that enable the high-fidelity, temporally accurate transmission required for sound localization. This highlights a critical role for developmental GABA co-release in shaping the specialized functional architecture of glycinergic synapses critical for binaural processing and sound localization.

## Introduction

Glycine and gamma-aminobutyric acid (GABA) are the two major inhibitory neurotransmitters in the mammalian central nervous system. In inhibitory synaptic terminals, both glycine and GABA are transported from the cytosol into synaptic vesicles by the shared vesicular GABA transporter (VGAT) (1, 2). Consequently, the relative cytosolic concentrations of glycine and GABA largely determine whether a neuron releases glycine, GABA, or both. The cytosolic concentration of glycine is determined primarily by the transmembrane glycine transporter 2, which is highly expressed in axonal boutons of glycinergic neurons (3) and transports glycine from the extracellular space into the synaptic cytosol. In contrast, GABA is synthesized directly in the cytosol from glutamate by glutamate decarboxylase (GAD), which exists in two isoforms, GAD65 and GAD67, encoded by independent genes (4).

Most inhibitory neurons release either glycine or GABA, but during development, many glycinergic synapses in the hindbrain, cerebellum, and spinal cord transiently express GAD and co-release GABA from synaptic vesicles that contain both glycine and GABA (5–7). Previous studies have shown that GABA release from mixed GABA/glycinergic synapses can affect the transmission at immature glycinergic synapses in multiple ways. For example, because GABA_A_ receptor-mediated currents decay more slowly than glycine receptor-mediated currents (5), activation of GABAA receptors at glycinergic synapses (8, 9) can prolong postsynaptic currents, leading to enhanced inhibition and temporal integration. On the other hand, because GABA can also act as a weak competitive agonist for glycine receptors (GlyRs), GABA co-release can shorten GlyR-mediated responses and reduce overall inhibition and temporal integration (10). Finally, co-released GABA can act on metabotropic GABA_B_ receptors, which, via activating G protein-coupled potassium channels, can increase and prolong postsynaptic inhibition or decrease presynaptic neurotransmitter release (11). However, despite increasing insights into the acute effects and synaptic mechanisms of GABA co-release, the longer-term role of GABA co-release from mixed glycinergic/GABAergic synapses in the maturation of glycinergic synapses and circuits remains poorly understood. At purely GABAergic synapses, GABA is important for the formation, maturation, and refinement of GABAergic synapses and circuits (12–14), and it is possible that by co-releasing GABA, developing glycinergic synapses and circuits gain access to these functions.

In the developing auditory brainstem and midbrain, GABA co-release from glycinergic neurons is widespread (7, 9, 15, 16). In the sound localization pathway from the medial nucleus of the trapezoid body (MNTB) to the lateral superior olive (LSO), glycinergic MNTB terminals co-release GABA and glutamate before the onset of hearing (in rodents approximately at end of the second postnatal week), a developmental period which coincides with the peak of synaptic maturation and tonotopic refinement via synaptic elimination and strengthening (17–21). During this refinement period, MNTB-LSO synapses exhibit activity-dependent long-term depression and potentiation, both of which depend on GABA receptor activation (22, 23), suggesting that GABA co-release participates in the synaptic mechanisms underlying the regulation of synaptic strength and tonotopic refinement.

To elucidate the roles of GABA co-release in the maturation and refinement of the MNTB-LSO pathway, we examined the synaptic and circuit organization of this pathway in conditional knockout mice, in which MNTB neurons lack the expression of both GAD isoforms and exhibit severely reduced co-release of GABA. Using whole-cell recordings in brainstem slices, we found that disrupting GABA co-release had no effect on the developmental elimination and strengthening of MNTB-LSO connections, indicating that these major refinement processes do not depend on GABA co-release. However, MNTB-LSO synapses in knockout mice exhibited a substantial increase in the quantal size and a significantly smaller readily releasable vesicle pool. These synaptic alterations negatively affected the fidelity and temporal precision of synaptic transmission, both of which are essential for accurately encoding binaural sound localization cues. Together, these results demonstrate that GABA co-release plays a critical role in the developmental acquisition of specialized functional architecture features of a central glycinergic auditory synapse.

## Results

### Genetic deletion of GAD65 and GAD67 impedes GABA co-release from MNTB-LSO synapses

The concentration of cytosolic GABA determines the concentration of vesicular GABA, and limiting cytosolic GABA supply interferes with synaptic GABA release (2, 24, 25). Therefore, to abolish GABA co-release from MNTB terminals, we blocked cytosolic GABA production in MNTB neurons by genetically deleting GAD65 and GAD67. To avoid the neonatal lethality associated with global GAD67 deletion (26), we took advantage of the fact that, in the brainstem auditory sound localization pathway, expression of the vesicular glutamate transporter 3 (vGlut3) is restricted to MNTB neurons (27, 28). We used vGlut3 promoter-induced Cre expression (29) to drive Cre-dependent excision of ‘floxed’ GAD67 in the MNTB. These conditional GAD67 knockout mice (Vglut3-IRES-Cre; GAD67^fl/fl^) were crossed with global GAD65 KO mice (65KO) to create viable GAD65/GAD67 double KO mice (dKO, Vglut3-IRES-Cre; GAD65^-/-^; GAD67^fl/fl^). Immunohistochemical labeling of GAD65 and GAD67 in the brainstem of wild-type mice (WT) (postnatal day P3 - 5) confirmed the expression of both GAD isoforms in MNTB axon terminals in the LSO, and the lack of GAD expression in dKO mice (Fig. 1). To verify that the deletion of both GAD isoforms disrupts GABA co-transmission at MNTB-LSO synapses, we recorded MNTB-elicited synaptic responses from LSO neurons in brainstem slices from 3-5 day-old mice (Fig. 1D-F), the age when GABA co-transmission at MNTB-LSO synapses is most predominant (6, 9, 30). Blocking glutamatergic AMPA receptors with CNQX (20 µM) and glycine receptors with strychnine (0.2 µM) isolated residual GABA-mediated currents (Fig. 1E). In WT mice, these GABAergic currents comprised 14% of MNTB-elicited currents, whereas in dKO mice, they were significantly reduced to only 4 % (Fig. 1F, p = 0.008, unpaired t-test), a 74% reduction compared to WT mice. Thus, deletion of GAD65 and GAD67 severely disrupts GABAergic co-transmission in the MNTB-LSO pathway, without affecting the overall response amplitude or significantly reducing the glycinergic component of MNTB-elicited responses (Fig. S1). The small, residual GABA-mediated component in dKO mice may reflect an incomplete block of GlyRs by 0.2 µM strychnine, GAD-independent GABA synthesis (31, 32), or presynaptic GABA uptake (33).

**Fig. 1.**
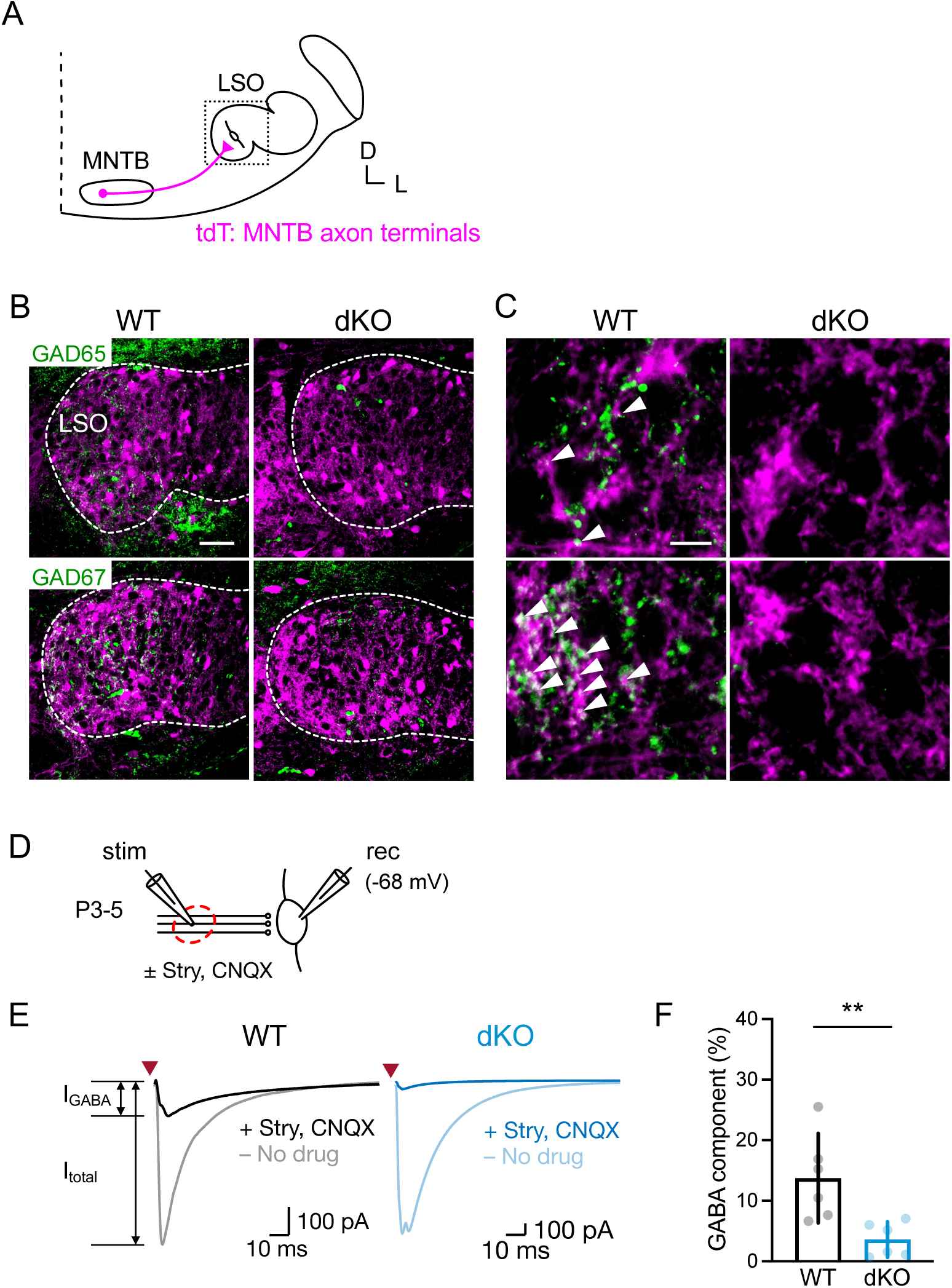
MNTB-LSO axon terminals in dKO mice lack the expression of GAD65 and GAD67 and have disrupted GABA co-transmission. **(A**) Schematic of the MNTB-LSO pathway in the mouse auditory brainstem. MNTB neurons provide a topographic projection to the LSO. MNTB axons are labeled by the expression of Vglut3 promoter-driven tdTomato (tdT, magenta). (**B**) Immunolabeling of GAD65 and GAD67 in the medial limb of the LSO from a WT (P5) and dKO mouse (P5). Confocal single optical sections. Dotted lines highlight the LSO boundary. Scale bar, 50 µm. **(C)** Higher magnification images. Arrowheads point to the co-labelling of GAD (green) and tdTomato (magenta). Scale bar, 10 µm. (**D**) Schematic illustration of whole-cell patch-clamp recordings from neurons in the LSO while stimulating the MNTB-LSO pathway. stim, stimulation electrode; rec, whole-cell recording electrode. **(E)** Example traces of MNTB-elicited postsynaptic currents from WT and dKO mice before and after application of strychnine (0.2 µM) and CNQX (20 µM). Red triangle indicates electrical stimulation. Stimulus artifacts are omitted for clarity. **(F)** The GABA component (CNQX/strychnine-insensitive current) in the dKO was residual and significantly smaller than in the WT (mean % of GABA component, WT vs dKO: 13.7 vs 3.6; unpaired t-test, *t*(*_10_*) = 3.27, *p* = 0.008).

### GABA-independent refinement of MNTB-LSO connectivity

During the first two postnatal weeks, the MNTB-LSO pathway undergoes activity-dependent refinement, including the synaptic silencing of most initially formed MNTB-LSO connections and the strengthening of those that are maintained (17, 19, 20). GABA co-release, possibly through its role in inducing long-term depression and long-term potentiation (23, 34, 35), or through its promotion of postsynaptic glycine receptor clustering (36), may play an important role in this refinement. To test this hypothesis, we characterized the number and strength of MNTB axons converging onto single LSO neurons at the end of the functional refinement period (P10-12) (17, 34). Electrical stimulation of the MNTB-LSO pathway with gradually increasing stimulation strengths (Fig. 2A) produced inhibitory postsynaptic currents (IPSCs) with increasing amplitudes due to the recruitment of an increasing number of MNTB axons (Fig. 2B). In some LSO neurons, recruitment of additional fibers resulted in a clear stepwise increase of IPSC amplitudes, but in most neurons, such steps were less noticeable. To determine the number and strength of individual axonal inputs, we analyzed the distribution of IPSC amplitudes using an unsupervised Gaussian mixture model (GMM), which assumes that the distribution of IPSC amplitudes elicited by multiple fibers reflects a linear combination of a Gaussian distribution of amplitudes from each fiber. Under this assumption, the peaks of each Gaussian fit to the population of peak amplitudes represent individual axonal inputs, and the distance between peaks represents their strength (Fig. 2B) (35, 36). For three genotypes (WT, 65KO, dKO), the number of input fibers and their average strengths were similar to what has been previously reported for mice (17, 18, 20, 30, 36) and, importantly, did not differ between genotypes (Fig. 2C-E). The strength of individual MNTB inputs varied widely (Fig. 2F-J) and was evenly distributed with no differences between genotypes. Because the estimation of input number and input strength could vary depending on the specific clustering approaches, we also analyzed IPSC amplitudes using k-means clustering and density-based spatial clustering of applications with noise (DBSCAN). As expected, these clustering approaches yielded slightly different estimates of the number and strength of fibers (Figs. S2, S3), but importantly, they also revealed no differences in the number of input fibers between genotypes. Together, these results indicate that the developmental circuit refinement of the glycinergic MNTB-LSO pathway is insensitive to a severe disruption of synaptic GABA co-release.

**Fig. 2.**
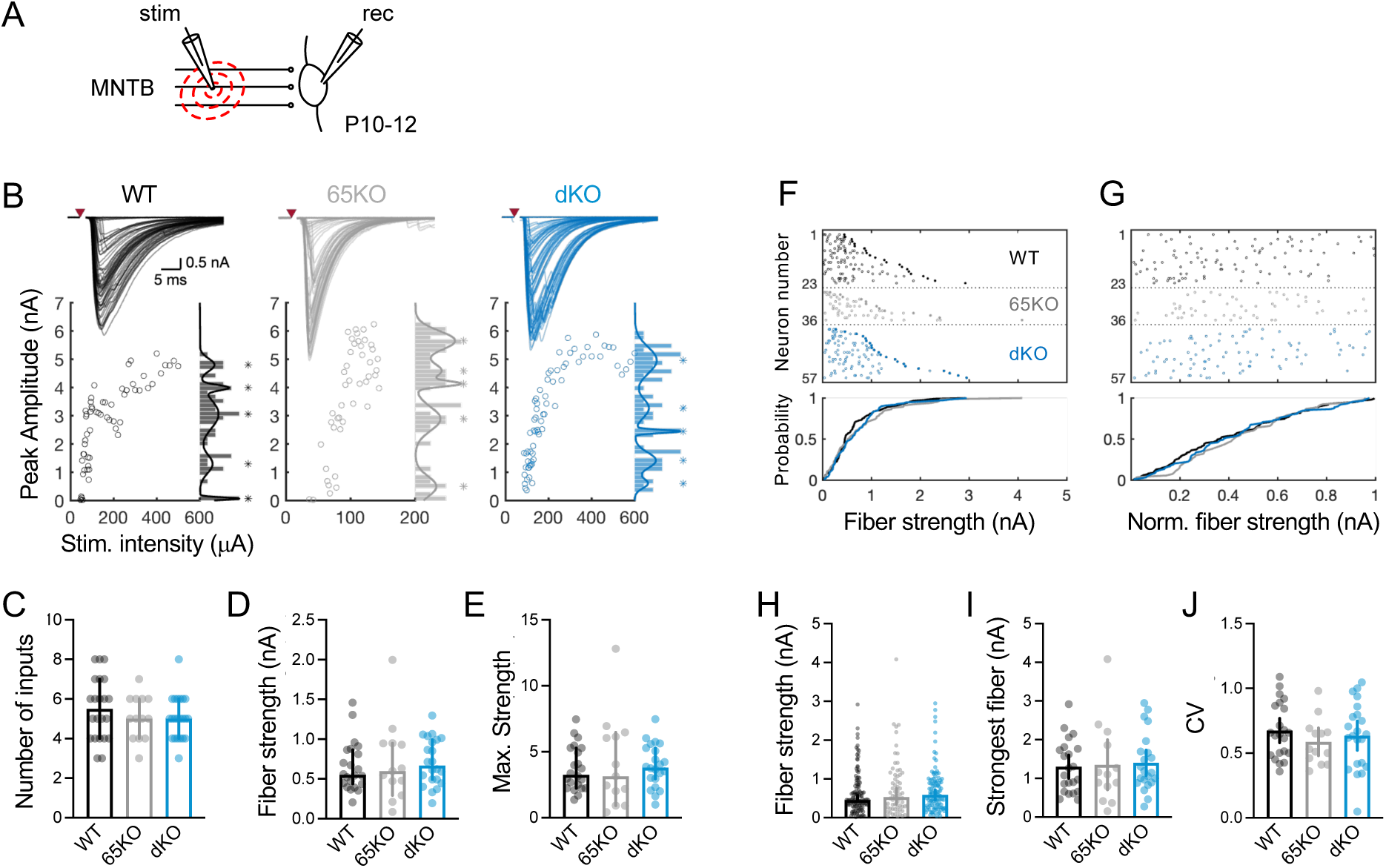
Lack of GABA co-release has no effect on the developmental pruning and strengthening of MNTB-LSO connections and does not influence fiber strength distribution. **(A)** To estimate the number and strength of MNTB inputs to individual LSO neurons, the MNTB fibers were stimulated by gradually increasing the stimulation current while recording from LSO neurons of mice, aged P10-P12. (B, *top*) Superimposed traces of the synaptic currents recorded in an example LSO neuron for each genotype (left to right: WT, 65KO, dKO). Red triangles indicate stimulation timing; stimulus artifacts are omitted for clarity. (B, *bottom*) Plots of peak amplitude versus stimulation intensity corresponding to the example neurons on top. Histograms (bin size, 0.15 nA) and multiple Gaussian fits on the right show the frequency distribution of peak amplitudes. Asterisks (*) indicate the center of individual Gaussian fits. The number of Gaussians indicates the number of fibers. The distance between the peaks of neighboring Gaussians indicates the fiber strength. **(C)** The number of input fibers, **(D)** the mean fiber strength, and **(E)** the maximum input amplitude in the population of LSO neurons was not different between genotypes (WT: n = 22, N = 15; 65KO: n = 13, N = 4; and dKO: n = 22, N = 10; number of input fibers: Kruskal-Wallis, *H*(*_2_*) = 1.11, *p* = 0.57; fiber strength: Kruskal-Wallis, *H*(*_2_*) = 0.91, *p* = 0.64; maximum input: One-way ANOVA, *F*(*_2, 54_*) = 0.18, *p* = 0.83). (F, *top*) Strength of all individual fibers for the same LSO neurons shown in (C-E). Filled circles indicate the strongest fiber of each neuron. (F, *bottom)* Fiber strength cumulative distribution superimposed for the three genotypes. The fiber strength distribution was not different between genotypes (Anderson-Darling K-sample test, *Anderson-Darling* = 1.95, *p* = 0.41). (**G**) Same data as in (F) but normalized to the strongest fiber for each neuron. The normalized fiber strength distribution was not different between genotypes (Anderson-Darling K-sample test, *Anderson-Darling* = 1.47, *p* = 0.58). The strongest fiber for each neuron is omitted from the top plot. (**H**) The median fiber strength when all individual fibers of all neurons were pulled together as a population was not different between genotypes (Median = 0.46, 0.53, and 0.59; n = 121, 66, and 111; for WT, 65KO and dKO, respectively; Kruskal-Wallis, *H_(2)_* = 1.15, *p* = 0.56). (**I, J**) The three genotypes did not differ in the CV of fiber strength (I, one-way ANOVA, *F_(2, 54)_* = 0.63, *p* = 0.53) and in the strength of their strongest fiber (J, one-way ANOVA, *F_(2, 54)_* = 0.09, *p* = 0.92).

### Disruption of GABA co-release impairs the development of the functional architecture of glycinergic MNTB-LSO synapses

The strength of a single-axon input is a function of the number of presynaptic vesicle release sites (N), the probability of a vesicle being released by an action potential (P_r_), and the magnitude of the postsynaptic response resulting from the release of neurotransmitter of a single vesicle (quantal size). Each of these parameters can be modulated by neuronal activity (37–40) and, in the MNTB-LSO pathway, undergo maturational changes (30, 41). We therefore investigated whether a disruption of GABA co-release alters the maturation of these synaptic parameters. To determine the quantal size of MNTB-LSO synapses, we blocked action potentials with tetrodotoxin (1 μM) and AMPA receptors with CNQX (20 µM) and recorded spontaneous miniature inhibitory postsynaptic currents (mIPSCs) (Fig. 3A, B), which closely represent the synaptic currents caused by the spontaneous release of single synaptic vesicles (42). We observed similar mIPSC amplitudes in WT and 65KO mice (Fig. 3B-D), which were also similar to what has been previously reported (1, 42). However, in dKO mice, the mean mIPSC amplitude was 67% larger than that in WT mice (median amplitude: WT, 38.4 pA; 65KO, 29.9 pA; dKO, 64.1 pA). In contrast, the frequency and the kinetics of mIPSCs were similar in all three genotypes. (Fig. 3E-G).

**Fig. 3.**
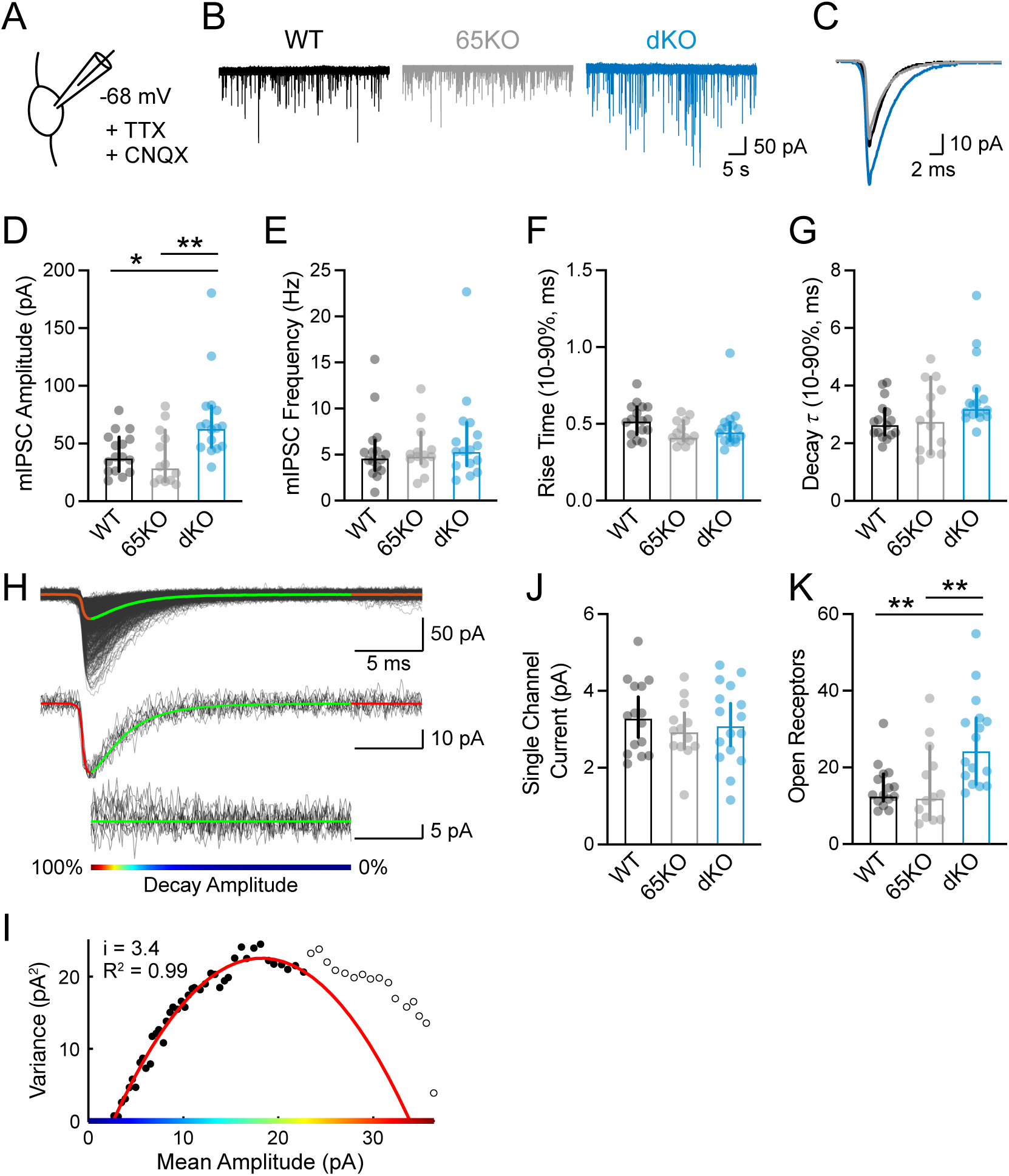
Disruption of GABA co-release increases the quantal size and number of postsynaptic receptors. **(A)** mIPSCs in LSO neurons were recorded in the presence of 1 µM TTX and 20 µM CNQX, at P10-12. **(B)** Example traces of mIPSCs from WT (black), 65KO (grey), and dKO (blue). **(C)** Superimposed, average traces of peak-aligned mIPSCs for neurons shown in (B). **(D)** mIPSCs amplitude, **(E)** frequency, **(F)** rise time, and **(G)** decay time values in the three genotypes (WT: n = 16, N = 6; 65KO: n = 13, N = 4; and dKO: n = 16, N = 6). The peak amplitude of mIPSCs was significantly increased in the dKO (D, Kruskal-Wallis, *H_(2)_* = 11.38, *p* = 0.003, Dunn’s test, WT vs dKO, *p* = 0.031, dKO vs 65KO, *p* = 0.005). There were no differences in mIPSC frequency (E, Kruskal-Wallis, *H_(2)_* = 0.97, *p* = 0.62), and kinetics (F, rise time: Kruskal-Wallis, *H_(2)_* = 4.46, *p* = 0.11; G, decay time: Kruskal-Wallis, *H_(2)_* = 0.64, *p* = 0.041, Dunn’s test, WT vs dKO, *p* = 0.052, dKO vs 65KO, *p* = 0.18). **(H-I)** estimation of single channel currents by peak-scaled non-stationary fluctuation analysis. **(H)** *Top:* Overlay of aligned mIPSCs from a neuron (WT, P10, 977 traces in black) and the mean mIPSC trace (red, decay in green). *Middle:* Mean mIPSC and 10 example mIPSCs with similar peak amplitudes (black). *Bottom:* Difference currents of mean trace (green) and individual traces (black) from mean decay. Color bar – percentage of decay amplitude over time. **(I)** Variance vs mean decay amplitude of the neuron shown in (H). The parabola was fitted to the first 75% of data (filled circles). The estimated single channel current (i) of this cell was 3.4 pA. **(J)** Estimated single current amplitudes were not different between genotypes. WT: 3.32 ± 0.52 pA, n = 15, N= 5; 65KO: 2.97 ± 0.47 pA, n = 13, N = 4; dKO: 3.12 ± 0.55 pA, n = 16, n = 6; One-way ANOVA, *F_(2, 41)_* = 0.51, *p* = 0.60. **(K)** The estimated number of open receptors (mean IPSC amplitude divided by mean single channel current) was significantly higher in dKO mice, compared to WT and 65KO mice. WT: 12.83 [11.22, 18.25]; 65KO: 12.29 [6.39, 25.50]; dKO: 24.64 [15.56, 32.85]; Kruskal-Wallis, *H_(2)_* = 13.05, *p* = 0.002; Dunn’s test, WT vs dKO, *p* = 0.006, dKO vs 65KO, *p* = 0.006). n, N same as in (J). \**p* < 0.05, \*\**p* < 0.01.

The increased mIPSC amplitudes in dKO mice could result from the activation of more postsynaptic glycine receptors, an increase in the conductance of individual receptors, or an increase in the amount of glycine released from individual synaptic vesicles. To estimate the mean single-channel current of individual glycine receptors, we analyzed mIPSCs using peak-scaled non-stationary fluctuation analysis (43)(SI Methods). For each neuron, mIPSCs were averaged, and the resulting mean mIPSC trace was peak-scaled to each individual trace (Fig. 3H). The variance of the individual traces against the mean trace during the decay was plotted against the mean amplitude and single-channel currents were derived from a parabola fit to the first 75% of data points. This analysis resulted in a mean single channel current for WT mice of 3.3 ± 0.52 pA (n = 15, N= 5), corresponding to a mean single channel conductance of 66.6 ± 10.6 pS, which is in the range of glycine receptor conductances reported previously (44, 45). Most importantly, the mean single channel current was very similar and statistically not different across all three genotypes (Fig. 3J), making it unlikely that the larger mIPSCs in dKO mice were caused by the synaptic expression of higher-conducting glycine receptors. Assuming that the quantal size of miniature evens and evoked events is similar, dividing the mean mIPSC amplitudes by the mean single channel current provides an estimate of the mean number of glycine receptors activated during an mIPSC. This estimation revealed 13 open glycine receptors in WT mice and 12 receptors in 65 KO mice, which was not statistically different (Fig. 3K). For dKO mice, however, the estimation was 25 open receptors, a 100% increase compared to WT and dKO mice. Although these estimates do not account for the possibility of an increased vesicular glycine content, our results indicate that a disruption of GABA co-release leads to a significantly increase in the number of postsynaptic glycine receptors at MNTB-LSO synapses.

The notable increase in quantal size despite unchanged strength of individual MNTB-LSO connections in dKO mice (Fig. 2D) was unexpected. This finding suggests that MNTB axons in dKO mice have fewer synaptic release sites (N), resulting in a smaller readily releasable pool (RRP), meaning fewer vesicles are available for immediate action potential-triggered release. To explore this possibility, we estimated the size of the RRP by stimulating the MNTB-LSO pathway with trains of 50 stimuli at 100 Hz and analyzing IPSCs using the train method (46) (Fig. 4). This method assumes that the vesicle replenishment rate is constant throughout the train stimulation and that stimulations reach the point at which vesicle release is balanced with replenishment. Therefore, in the cumulative amplitude plot, the intercept of a regression line fitted to the last five points of the cumulative IPSC with the ordinate denotes the RRP (the cumulative amount of release minus the release by replenished vesicles). The slope of the regression line estimates the vesicle replenishment rate. This analysis revealed that the RRP in dKO mice was reduced to 57% of the RRP of WT or 65KO mice (Fig. 4D). The estimated replenishment rate, normalized to the size of the RRP, was not different between genotypes (Fig. 4E). The probability of release (P_r_), calculated from the ratio of the amplitude of the first IPSC and the RRP, was low (∼ 0.15) and indistinguishable between WT and dKO mice, although we observed a slight decrease of P_r_ between 65KO and dKO mice (Fig. 4F). These results indicate that in dKO mice, a 67% increase in quantal size (Fig. 3D) is compensated by a 43% decrease in the RRP (Fig. 4D), leading to unchanged single-axon strength (Fig. 2D).

**Fig. 4.**
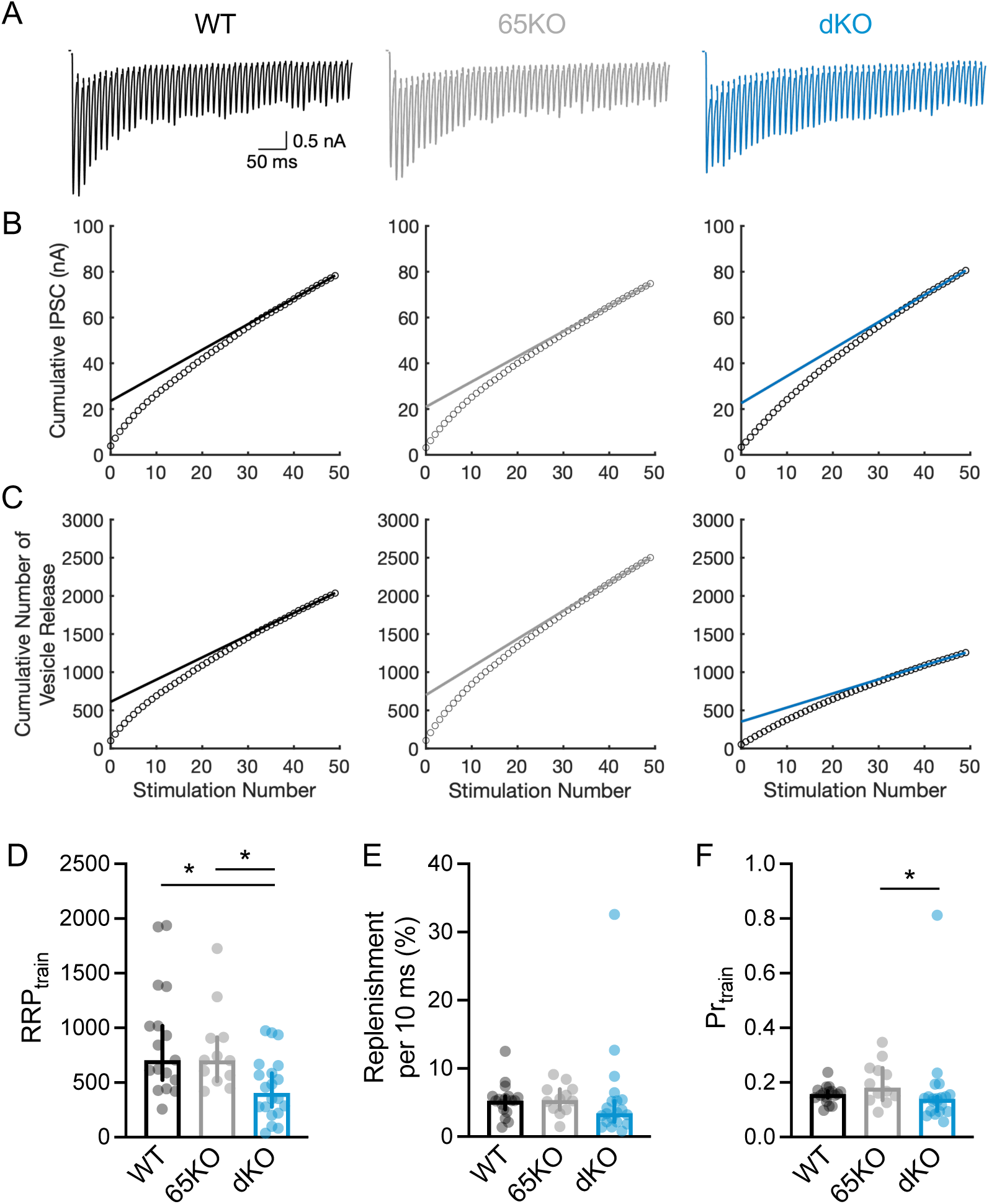
Lack of developmental GABA co-release reduces the size of the readily releasable pool of vesicles. **(A)** Example IPSC traces recorded in response to a train of 50 stimuli at 100 Hz in WT, 65KO, and dKO mice at age P10-14. Stimulus artifacts are omitted for clarity. **(B)** Examples of the estimation of the RRP size in the neurons represented in (A) by using the train method(46). The relative size of the RRP corresponds to the y-axis intercept of a linear regression fit back-extrapolated from the last five points of cumulative IPSC amplitudes obtained by train stimulation (open circles). **(C)** Cumulative number of released vesicles as calculated by dividing IPSC amplitudes by quantal size (WT, 38.45 pA; 65KO, 29.91 pA; dKO, 64.09 pA). **(D)** The RRP in dKO mice was significantly reduced as compared to WT and 65KO mice (Kruskal-Wallis, *H_(2)_* = 10.37, *p* = 0.006, Dunn’s test, WT vs dKO, *p* = 0.010, dKO vs 65KO, *p* = 0.044). **(E)** The replenishment rate normalized to the RRP was not different between genotypes (Kruskal-Wallis, *H*(*_2_*) = 4.06, *p* = 0.13). **(F)** Vesicle release probability in dKO mice was not different from WT mice but was lower than in 65KO mice (Kruskal-Wallis, *H_(2)_* = 6.73, *p* = 0.037, Dunn’s test, WT vs. dKO, *p* = 0.38, dKO vs. 65KO, *p* = 0.033). \**p* < 0.05.

### Disruption of developmental GABA co-release impairs the high-fidelity of synaptic MNTB-LSO transmission

A hallmark of MNTB-LSO connections is their very large quantal content and a low P_r,_ which are thought to underly their remarkably high level of fidelity (low failure rates of eliciting postsynaptic responses) even during high, prolonged activity (47, 48). Despite exhibiting a normal strength in response to single stimuli, the lower quantal content of MNTB-LSO connections in dKO mice could thus reduce synaptic fidelity during prolonged activity. To investigate this possibility, we examined the success rate of synaptic transmission of MNTB-LSO connections during 60-second-long stimulation trains at 50 Hz (Fig. 5). In both WT and 65KO mice, MNTB-LSO connections maintained a fidelity rate of over 70% even after 2500 stimulations, similar to what has been previously described (47, 48) (Fig. 5C, D). In contrast, in dKO mice, synaptic fidelity decreased to less than 40%. This decrease is unlikely to result from a failure to activate all presynaptic axons, because in dKO mice, fidelity stayed near 100% during the first 150 stimuli (Fig. S5). Additionally, the time until fidelity began to decline (latency, Fig. 5B,E) was significantly shorter in dKO mice compared to WT or 65KO mice, and the rate of decline was significantly faster (Fig. 5F). In addition to lower synaptic fidelity, synapses in dKO mice also exhibited faster and more pronounced synaptic depression, as indicated by smaller IPSC amplitudes during the first 2-3 seconds and the last 10 seconds of the train (Fig. 5G-I). When synaptic failures were excluded, dKO mice still showed more synaptic depression at the beginning of the train (Fig. 5J-L, S2), suggesting that the increased synaptic depression in dKO mice at the steady state level primarily reflects an increase in failure rate.

**Fig. 5.**
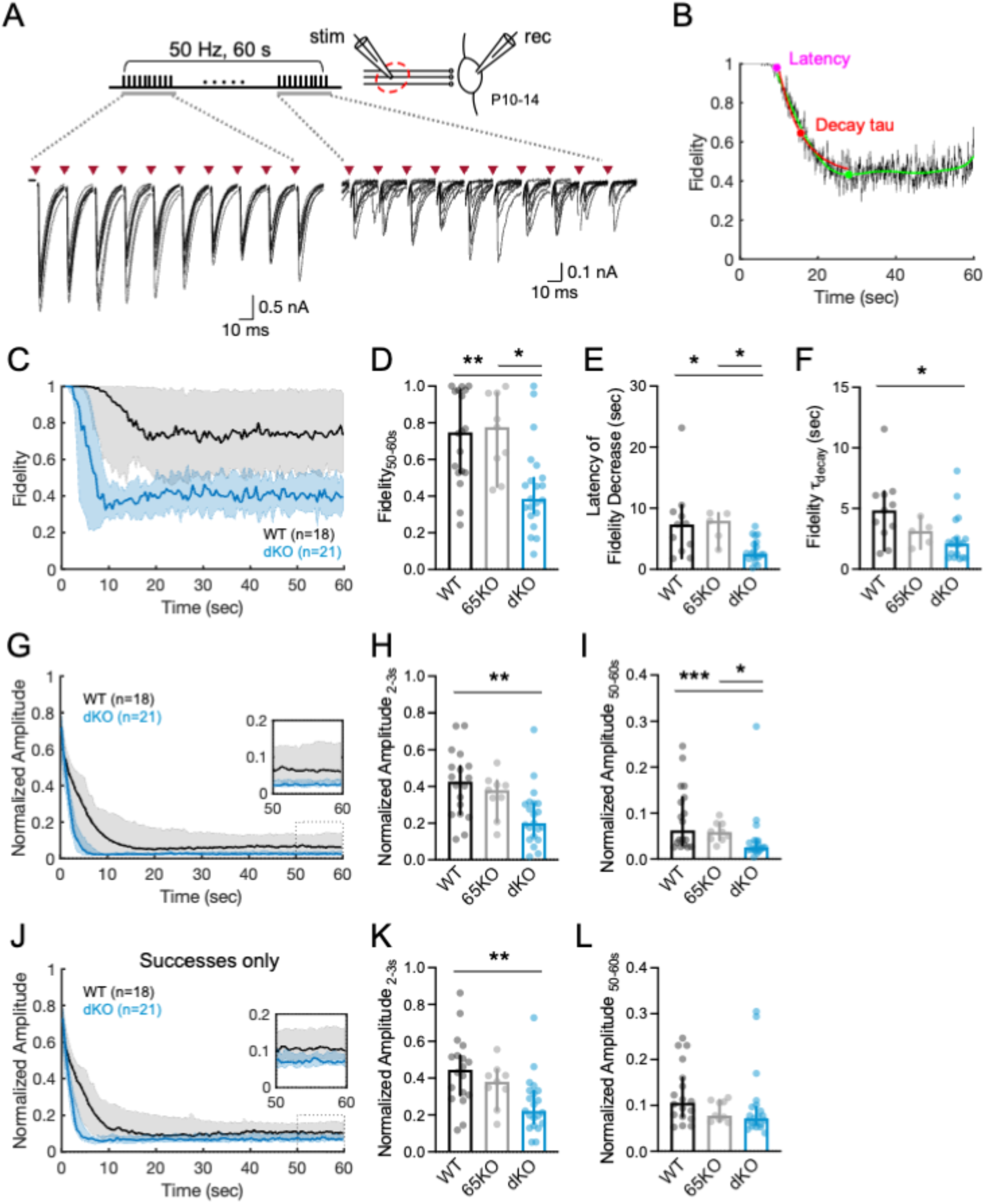
Disruption of GABA co-release impairs synaptic fidelity during prolonged stimulations. **(A)** Experimental design. MNTB fibers were stimulated at 50 Hz for 60 s. Example traces (9 iterations) from a WT neuron (P12) illustrating the postsynaptic responses evoked by the first 10 (left) and last 10 (right) stimuli. stim, stimulation electrode on MNTB fibers; rec, recording electrode. Red triangles indicate the stimulation. Stimulation artifacts are omitted for clarity. **(B)** Fidelity (rate of successful responses) of the example neuron in (A). Fidelity is plotted as a moving average of 10 data points. For quantitative analyses, fidelity data were force-fitted with the 8^th^ order of a polynomial equation (green line) from the time when fidelity begins to drop (latency, magenta circle). The latency and the time of the first valley derived from the polynomial fit (green circle) were then used as the start and end time points to fit a single exponential to the data (red line). The single exponential was then used to derive the decay tau (red circle). **(C)** Population plot of fidelity (median ± 95% CI, bin size 0.5 s) for WT and dKO mice. For clarity, data from 65KO mice are omitted. **(D)** Fidelity during the last 10 s of the train was significantly lower in dKO mice than in WT or 65KO mice (Kruskal-Wallis, *H_(2)_* = 12.79, *p* = 0.002; Dunn’s test, WT vs dKO, *p* = 0.004, WT vs 65KO, *p* > 0.99, dKO vs 65KO, *p* = 0.023). (WT: n = 18, N = 8; 65KO: n = 9, N = 6; dKO: n = 21, N = 6). **(E)** Latency to drop of fidelity was significantly shorter in dKO mice than in WT or 65KO mice (Kruskal-Wallis, *H_(2)_* = 10.19, *p* = 0.006; Dunn’s test, WT vs dKO, *p* = 0.030, WT vs 65KO, *p* > 0.99, dKO vs 65KO, *p* = 0.035). **(F)** Fidelity decrease (tau) was significantly faster in dKO mice than in WT mice (Kruskal-Wallis, *H_(2)_* = 8.02, *p* = 0.018; Dunn’s test, WT vs dKO, *p* = 0.015, WT vs 65KO, *p* = 0.95, dKO vs 65KO, *p* = 0.88). In (E) and (F), neurons with a latency longer than 55 s were excluded (WT, n = 7 out of 18; 65KO, n = 4 out of 9; dKO, n = 2 out of 21). **(G)** Population plot of IPSC amplitudes normalized to the first response (median ± 95% CI bin size, 0.5 s). **(H)** Normalized IPSC amplitudes in dKO mice are significantly reduced 2-3 s after train onset compared to WT mice (Kruskal-Wallis, *H_(2)_* = 11.37, *p* = 0.003; Dunn’s test, WT vs dKO, *p* = 0.004, WT vs 65KO, *p* > 0.99, dKO vs 65KO, *p* = 0.11). **(I)** Steady state depression in dKO mice was stronger than in WT and 65KO mice (Kruskal-Wallis, *H_(2)_* = 15.69, *p* < 0.001; Dunn’s test, WT vs dKO, *p* < 0.001, WT vs 65KO, *p* > 0.99, dKO vs 65KO, *p* = 0.013). **(J)** Normalized IPSC amplitudes of successful responses only (bin size, 0.5 s). **(K)** Normalized amplitudes in dKO mice are significantly reduced during 2-3 s after train onset (Kruskal-Wallis, *H_(2)_* = 11.35, *p* = 0.003; Dunn’s test, WT vs dKO, *p* = 0.003, WT vs 65KO, *p* > 0.99, dKO vs 65KO, *p* = 0.13). **(L)** Steady state depression was indistinguishable between genotypes (Kruskal-Wallis, *H_(2)_* = 4.90, *p* = 0.09). \**p* < 0.05, \*\**p* < 0.01, \*\*\**p* < 0.001.

The large quantal content of MNTB-LSO connections also underlies their remarkable temporal precision, which is crucial for the accurate encoding of binaural sound localization cues by LSO neurons (47, 49, 50). The lower quantal content of MNTB-LSO connections in dKO mice predicts larger trial-to-trial fluctuations of the mean latency of vesicle release during prolonged high activity, thereby broadening the temporal jitter of the peak of evoked IPSCs (Fig. 6A). To test this prediction, we analyzed the standard deviation of the latency of the IPSC peaks (SD_Peak Latency_) during train stimulation (Fig. 6B). The SD_Peak Latency_ rapidly increased during the first few seconds of a train before stabilizing in a plateau phase. For 50-Hz stimulations, we found no difference in the SD_Peak Latency_ between dKO mice and WT or 65KO mice (Fig. 6C, E, F). However, when challenged with 100 Hz trains, the SD_Peak Latency_ during the plateau phase was significantly increased in dKO mice (Fig. 6D, G, H). The increased jitter in dKO mice raises the possibility that some delayed responses escaped detection in our analysis, which only included responses with a peak latency < 5 ms to avoid inclusion of possible polysynaptic responses. The exclusion of these longer-latency responses could have resulted in an underestimation of the fidelity rate and jitter. Indeed, inclusion of longer-latency responses (up to 15 ms for 50 Hz stimulation and up to 8 ms for 100 Hz stimulation) slightly increased fidelity (Fig. S4) and jitter (Fig. S6), but it did not change the observed differences between genotypes. Together, these results indicate that disrupted GABA co-transmission during the development of MNTB-LSO connections reduces their temporal precision at physiologically relevant activity levels and strongly supports the idea that their large RRP size is a mechanism to attain temporally precise synaptic transmission.

**Fig. 6.**
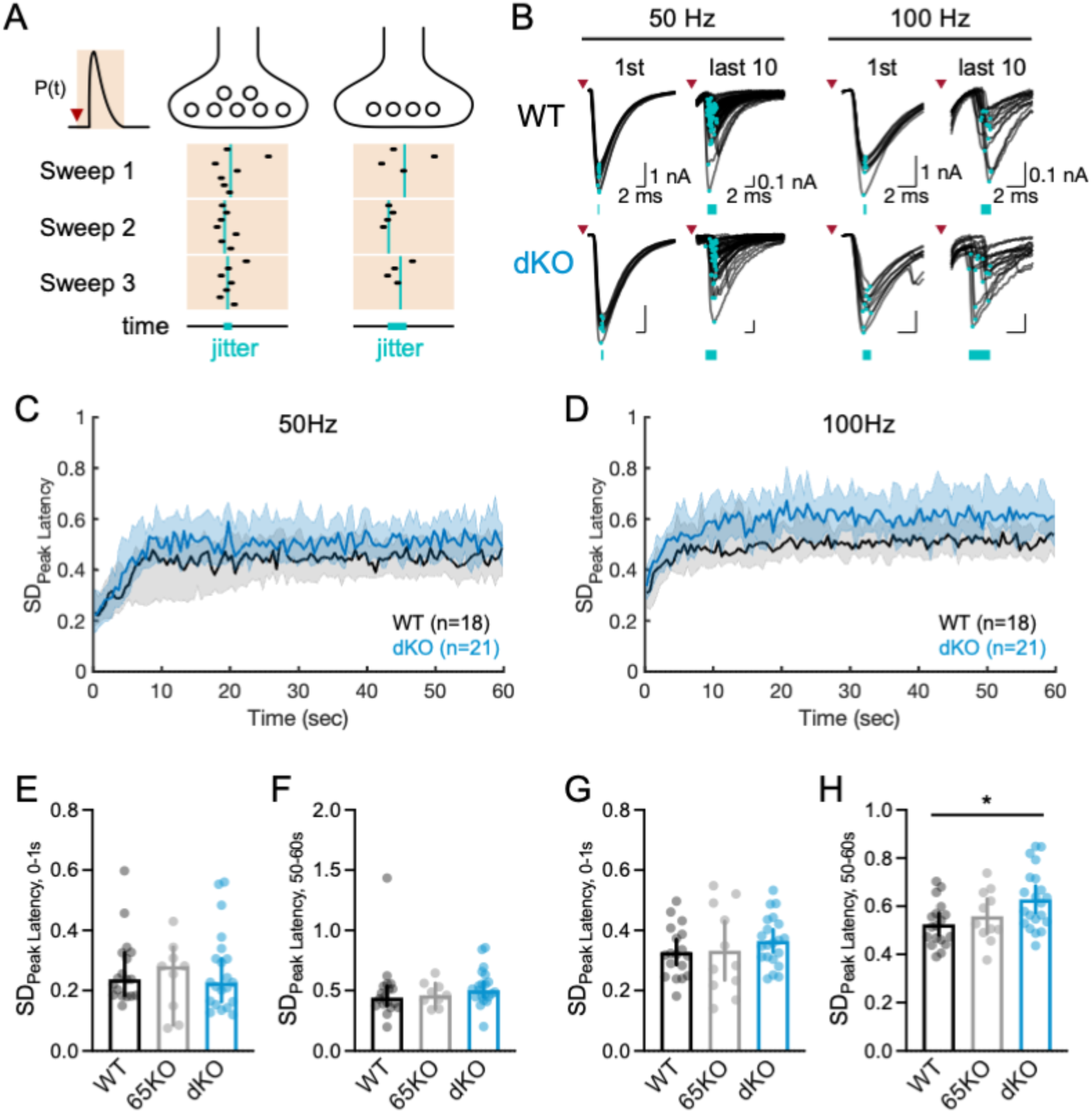
Lack of GABA co-release decreases temporal precision of synaptic transmission under high-frequency activity. **(A)** Scheme illustrating the relationship of RRP size with the temporal precision of synaptic transmission (see (47). P(t) illustrates the release probability for a single vesicle after electrical stimulation (red triangle). Black dots represent single vesicle release, and turquoise lines represent the mean latency of released vesicles. The smaller RRP in dKO mice may increase the temporal jitter of the mean latency of released vesicles, thereby lowering the temporal precision. **(B)** Superimposed traces of successful postsynaptic responses to the first and last ten stimulations at 50-Hz (left) and 100-Hz (right) in an example LSO neuron from a WT mouse (top) and a dKO mouse (bottom). Peaks are marked by turquoise dots, and their jitter is indicated by turquoise bars. **(C)** Standard deviation of peak latency (median and 95% CI) throughout stimulus trains (0.5 s bins) at 50 Hz, and **(D)** at 100 Hz in WT and dKO mice. To account for fewer successful events in dKO mice (Fig. 5C, D), SD_Peak Latency_ was computed from 100 bootstrapped successful responses in each 0.5-s bin. For clarity, 65KO data are not shown in (C) and (D). For 100-Hz trains, the increase in in SD_Peak Latency_ was signififcnaly larger in dKO mice (Two-way RM ANOVA, *F_(13.1, 614.0)_* = 14.21, *p* < 0.001, time; *F_(2, 47)_* = 3.73, *p* = 0.031, genotype; *F_(238, 5593)_* = 0.89, *p* = 0.88, interaction) (WT: n = 18, N = 8; 65KO: n = 11, N = 7; dKO: n = 21, N = 6). **(E)** SD_Peak Latency_ in the beginning (0-1 s), and (**F**) at the end (50-60 s) of 50-Hz trains. At 50 Hz, SD_Peak Latency_ did not differ between genotypes (0-1 s: Kruskal-Wallis, *H_(2)_* = 0.24, *p* = 0.89; 50-60 s: Kruskal-Wallis, *H_(2)_* = 2.90, *p* = 0.23). (**G, H**) Same as (E, F) but for 100 Hz trains. At the end of 100 Hz trains, SD_Peak Latency_ was significantly larger in dKO mice than in WT mice (one-way ANOVA, *F_(2, 47)_* = 4.49, *p* = 0.016; Tukey’s test, WT vs dKO, *p* = 0.014, WT vs 65KO, *p* = 0.71, dKO vs 65KO, *p* = 0.21). \**p* < 0.05.

## Discussion

The widespread co-release of GABA from developing glycinergic synapses in the brain and spinal cord has been extensively documented (5, 8). But despite increasing insights into the acute effects of GABA co-release on synaptic transmission, the specific roles of this co-release in the development of glycinergic synapses or circuits have remained elusive. In the rodent primary sound localization circuit, glycinergic MNTB-LSO connections transiently co-release GABA during the first two postnatal weeks (6, 9, 30), coinciding with the period of synaptic maturation and circuit reorganization (17–21). The present results from mice with a disrupted GABA co-release from MNTB-LSO synapses suggest that GABA co-transmission is not required for the developmental pruning and strengthening of MNTB-LSO connections. However, disruption of GABA co-release impairs the maturation of the specialized synaptic architecture that enables high-fidelity and precisely timed transmission during sustained activity. These findings reveal a previously unrecognized developmental role for GABA co-release in shaping the functional properties of glycinergic circuits.

In the MNTB-LSO pathway, GABA co-release occurs during a period when most initially formed MNTB-LSO connections are silenced, and the remaining ones are strengthened, leading to an increase in tonotopic precision. (19, 20). This circuit remodeling is activity-dependent and contingent on the precise temporal pattern of spike activity that originates in the pre-hearing cochlea (17, 51). While the mechanisms by which patterned activity guides the tonotopic sharpening of the MNTB-LSO pathway are incompletely understood, before hearing onset, MNTB-LSO synapses can express LTP and LTD, both of which require the activation of postsynaptic GABA receptors (22, 23, 52). Because LTP and LTD are considered early steps in activity-dependent circuit refinement (53), we anticipated that disrupted GABA co-release would interfere with the silencing and/or strengthening of MNTB-LSO connections, leading to an abnormal strength or number of axonal inputs. This, however, was not the case as both the number as well as the strength of MNTB inputs to LSO neurons were unaffected in dKO mice (Fig. 2). These results could indicate that GABA-mediated LTP and LTD at MNTB-LSO synapses serve roles different from circuit refinement or that the disruption of GABA transmission in dKO mice leads to the expression of novel, hitherto unidentified, forms of LTP and LTD at MNTB-LSO synapses that do not rely on GABA release.

Although the connectivity and strength of MNTB-LSO connections remained unchanged in dKO mice, the disrupted GABA co-release compromised the development of the functional synaptic architecture of MNTB-LSO synapses. In dKO mice, the quantal size was almost 70% larger than in WT mice, and this increase correlated with the activation of twice as many postsynaptic glycine receptors. This estimation is based on the assumption that changes in quantal size reflect in glycine receptor number or properties, which is supported by previous studies, which demonstrated that quantal size correlates with the number of postsynaptic GlyRs (54, 55). However, our experiments cannot rule out that an increase in vesicular glycine content also contributed to the larger quantal size in dKO mice. The observed two-fold increase in glycine receptor number may therefore represent an overestimate. GABA and glycine compete for VGAT transport (24, 56), and a lower cytosolic GABA concentration in dKO mice may thus lead to a higher vesicular glycine concentration. However, such an increase is expected to be small because even at the developmental peak of GABA co-transmission, GABA-mediated currents account for only 14% of the total synaptic current (Fig. 1). In accordance with this, the magnitude of evoked glycinergic currents was similar in WT and dKO mice (Fig. S1). Furthermore, the developmental downregulation of GAD expression and GABA co-release from MNTB synapses is largely complete around hearing onset (6, 57), suggesting that any differences in the cytosolic GABA concentration between WT and dKO will be minor at the age at which we performed our recordings. Thus, while future experiments will be required to determine the extent to which a higher vesicular glycine may have contributed to the increased quantal size in dKO mice, its contribution is likely to be minor.

Despite a significant increase in the quantal size in dKO mice, the strength of individual MNTB-LSO connections remained the same, indicating that in dKO mice, evoked IPSCs were generated by fewer released vesicles. This could reflect a lower P_r_ or fewer synaptic release sites. Because in dKO mice, the P_r_ was unchanged, but the RRP was significantly reduced, MNTB-LSO connection most likely had fewer release sites. The precision with which single-fiber strength was maintained is remarkable, suggesting that the strength of MNTB-LSO connections is tightly controlled by homeostatic mechanisms. In the auditory system, activity-dependent homeostatic regulation of synaptic transmission occurs already at early stages of the auditory processing and can involve changes in RRP, P_r_, and quantal size (37, 40, 58, 59). Most of these studies focused on glutamatergic synapses, much less is known about the homeostatic mechanisms of glycinergic auditory synapses. Our results add to our knowledge about these mechanisms by demonstrating homeostatic regulation of quantal size and RRP at MNTB-LSO connections. The initiation trigger and the temporal sequence of this homeostatic regulation are currently unknown. However, due to GABA’s powerful effects on promoting synapse formation and stabilization (14, 60), it is conceivable that a disruption of GABA co-release impedes the formation and/or the stabilization of glycinergic MNTB-LSO synapses, resulting in connections with fewer release sites. The reduced strength of these connections could then trigger the homeostatic addition of postsynaptic GlyRs at existing synapses. This scenario is supported by previous findings demonstrating that the homeostatic regulation of inhibitory strength at mixed GABA/glycinergic synapses on spinal neurons is achieved by a postsynaptic trapping of GlyRs (55). At these synapses, the addition of glycine receptors is controlled by the activity of glutamatergic synapses and the activation of N-methyl-D-aspartate receptors (NMDARs). A similar NMDAR-dependency of glycinergic plasticity has also been reported for the activity-dependent postsynaptic potentiation of MNTB connections to the medial superior olive, which requires coincident activation of converging glutamatergic and glycinergic inputs (61). Interestingly, before hearing onset, NMDAR-dependent recruitment of synaptic GlyRs at MNTB-LSO synapses may not require the activation of glutamatergic inputs but could occur in a synapse-autonomous manner, because developing MNTB neurons express vGlut3 and co-release glutamate, which can activate postsynaptic NMDARs (28, 62). This scenario is supported by previous findings showing that a loss of glutamate co-release from developing MNTB-LSO synapses resulted in a smaller glycinergic quantal size (18).

To accurately encode the azimuthal direction of incoming sound, the synaptic inputs to LSO neurons must be highly secure and precisely timed (49). Furthermore, these properties must be maintained at high, sustained activity levels, as MNTB neurons *in vivo* exhibit spontaneous discharge rates up to 100 Hz and sound-evoked rates that can exceed 400 Hz (48, 63). To meet these challenges, MNTB-LSO connections possess a very large RRP, high rates of vesicle replenishment, and a low P_r_ (47). Our results from WT mice are consistent with these previous findings. The decreased RRP and synaptic fidelity in dKO mice (Fig. 5) provide experimental support for a crucial role of a large RRP of MNTB-LSO connections in enabling secure synaptic inhibition at natural activity levels. Similarly, the increased latency jitter observed in dKO mice substantiates the idea that a large RRP size is also critical for achieving high temporal precision. The impaired fidelity and temporal precision in dKO mice predict diminished sound localization accuracy in these animals. However, due to the high seizure and mortality rates of dKO mice during the third postnatal week (unpublished observation), likely caused by a disruption of GABA release from hippocampal and neocortical interneurons (64), testing these predictions will require MNTB-specific mouse driver lines.

In summary, our results from the developing auditory MNTB-LSO pathway demonstrate that GABA co-release from glycinergic synapses is not required for developmental circuit refinement through synaptic elimination and strengthening. Instead, GABA co-release is essential for establishing the specialized synaptic architecture of MNTB-LSO connection by promoting the formation of numerous weak synapses, which together create the large readily releasable pool (RRP) that ensures the high reliability and temporal precision of synaptic transmission that underly accurate sound localization.

## Materials and Methods

Additional description of methods can be found in *SI Appendix*.

### Animals

Mice of both sexes between postnatal day 2 (P2) and P14 were used. Double knockout mice for conditional deletion of GAD67 and germline deletion of GAD65 were produced by first crossing Vglut3-IRES-Cre mice (gift from Dr. Bradford Lowell, Harvard University) with GAD67^fl/fl^ mice (gift from Dr. Richard Palmiter, University of Washington) and resulting homozygote Vglut3-Cre; GAD67^fl/fl^ offspring were crossed with GAD65^+/–^ mice (strain # 003654, Jackson Laboratory, ME). Vglut3-Cre^+^; GAD67^f/lfl^; GAD65^−/−^ offsprings were used for dKO mice in experiments. For labeling MNTB neurons with tdTomato (tdT), Vglut3-Cre; GAD67^f/lfl^; GAD65^−/−^ were crossed with Ai9 reporter mice (strain #007909, Jackson Laboratory, ME) and Vglut3-Cre^+^; GAD67^f/lfl^; GAD65^−/−^; tdT^+^ were used for experiments. GAD67^fl/fl^, GAD67^fl/fl^; tdT, or tdT mice were used as WT controls. All experimental procedures were conducted in accordance with National Institute of Health guidelines and were approved by the Institutional Animal Care and Use Committee at the University of Pittsburgh.

### Immunohistochemistry

GAD65 and GAD67 were visualized in 50 μm coronal brainstem sections using antibodies for GAD65 (GAD-6, developed by D.I. Gottlieb and obtained from the Developmental Studies Hybridoma Bank, created by the NICHD of the NIH and maintained at The University of Iowa, Department of Biology, Iowa City, IA 52242 or GAD67 (MAB5406, MilliporeSigma, MA). Visualization of tdT expression was enhanced by immunolabeling using an anti-tdT antibody (AB8181, Sicgen). The secondary antibodies were conjugated to AlexaFluor-488 or AlexaFluor-568 (Thermo Fisher Scientific, MA). Confocal imaging was performed with LSM 700 (Zeiss, Germany) and analyzed with ZEN software (Zeiss, Germany).

### Electrophysiological slice recordings

Coronal brainstem slices (300 µm) were prepared as previously described (35). For recording, slices were bathed in recirculating ACSF (in mM: 124 NaCl, 1.3 MgSO_4_, 5 KCl, 1.25 KH_2_PO_4_, 10 Dextrose, 26 NaHCO_3_, 2 CaCl_2_. pH 7.4) at 22-25 ℃ (for experiments in Figs. 2 and 3) or 28 ℃ (for experiments in Figs. 4-6). Whole-cell patch-clamp recordings in voltage clamp were performed with pipettes (3-7 MΩ) filled with internal solution (in mM: 67.6 D-gluconic acid, 49 CsOH, 56 CsCl, 1 MgCl_2_, 1 CaCl_2_, 10 HEPES, 11 EGTA, 2 Mg-ATP, 0.3 Na-GTP, 3 Na_2_-phosphocreatine, 5 mM QX-314, adjusted to pH 7.2 and 280 mOsm. Membrane currents were digitized (10 kHz sampling rate) and filtered (3 KHz Bessel) with a MultiClamp 700B amplifier controlled by Clampex 10.7 software. Access resistance was compensated by 50% and membrane voltage was correct for the calculated junction potential (7.8 mV). Only LSO principal neurons characterized by the presence of a hyperpolarization-activated inward current (> 50 pA by hyperpolarization from -67.8 mV to -107.8 mV) were included in this study. Miniature IPSCs were recorded while blocking action potentials with tetrodotoxin (TTX, 1 mM) and AMPA receptors with CNQX (20 µM). mIPSCs were automatically detected with MiniAnalysis (Synaptosoft), followed by visual inspection.

### Estimation of MNTB fiber inputs

To estimate the number of MNTB axons converging on single LSO neurons, the three clustering methods Gaussian mixture model, k-means, and density-based spatial clustering of applications with noise (DBSCAN) were applied using MATLAB functions (MathWorks). Detailed descriptions are provided in *SI Appendix*.

### Peak-scaled nonstationary fluctuation analysis

To estimate single channel currents, mIPSCs were analyzed following published procedures (43). A detailed description is provided in *SI Appendix*

### Analysis of fidelity and temporal accuracy of postsynaptic responses during stimulation trains

Postsynaptic responses during train stimulation at maximum intensity were detected as peaks exceeding baseline + 3 SD within a post-stimulus window of 1.5 - 5.5 ms (Figs. 5 and 6) or 1.5-15 ms (50 Hz trains) or 1.5 1.5-8 ms (100 Hz trains) (Figs. S3 and S4). Fidelity was defined as the proportion of stimuli that elicited a detectable PSP, and for each neuron was calculated from 5-10 train repetitions. To derive the kinetics of fidelity decline, the fidelity of each neuron was plotted as a function of time using a moving average of 10 data points (Fig. 5B). The latency of fidelity decrease was determined as the time point at which the fidelity permanently dropped below 0.98. To determine the tau of the fidelity decline, an 8th-order polynomial force-fitted to the fidelity curve was used to find the first fidelity valley and a single exponential was then fitted to the fidelity curve from the latency to the first valley to calculate tau.

Temporal precision of postsynaptic responses during stimulation trains was determined as the variation in peak latencies of responses relative to the stimulation. To account for different numbers of responses across time and between genotypes, the standard deviation of peak latencies was computed using 100 bootstrapped successful responses in each 0.5-second bin.

### Statistical analyses

Data normality was assessed using the Shapiro-Wilks test using Prism (GraphPad, CA). Descriptive statistics are reported as mean ± 95% confidence interval (CI) for normally distributed data, and median with lower and upper bounds of 95% CI for non-normally distributed data. Statistical analyses to compare between groups were conducted using appropriate tests and post-hoc analyses as indicated in the text using Prism (GraphPad, CA) or MATLAB (Mathworks, MA) (64). The number of observations is denoted as “n” for the number of cells and “N” for the number of animals.

## Acknowledgments

We thank Dr. Richard Palmiter, University of Washington, and Dr. Bradford Lowell, Harvard University, for sharing mouse lines and Dr. Dong Hoon Shin and Dr. Juhoon So, University of Pittsburgh, for sharing equipment and for help with confocal imaging. We are grateful to Dr. Srivatsun Sadagopan for providing valuable support for the implementation of the Gaussian mixture model and to Dr. Flora Antunes for discussions and critical feedback on the manuscript. This work was supported by grants (5R01DC004199, 5R01DC019814) from the National Institute on Deafness and Other Communication Disorders.

## Supporting Information for

### Material and Methods

#### Animal genotyping

Genotyping was performed by polymerase chain reaction (PCR) from genomic DNA extracted from tail tissues with primer sequences (Table S1).

#### Immunohistochemistry

Mice at P5 were deeply anesthetized with isoflurane and transcardially perfused with PBS followed by 4% paraformaldehyde (PFA). The brains were removed, postfixed in 4% PFA for 2 days and cryoprotected by immersion in 30% sucrose for at least 2 days. The brain was blocked and coronal sections at 50 µm thickness were cut from the brainstem using a freezing microtome (HM 430, Epredia, MI). Sections were washed in PBS and incubated in 5% bovine serum albumin (in PBS) solution and M.O.M blocking reagent (Vector Laboratories, CA). Sections were then incubated in primary antibodies against either GAD65 (1:100, GAD-6, DSHB, IA), GAD67 (1:500, MAB5406, MilliporeSigma, MA), or tdT (1:500, AB8181, Sicgen) for 2 days at 4 ℃. Sections were washed three times in PBS and incubated with secondary antibody goat anti-mouse conjugated to AlexaFluor-488 or AlexaFluor-568 (1:1000, Thermo Fisher Scientific, MA), for 2 hours at 22-25 ℃. Sections were washed three times in PBS, mounted on coated slides, and cover-slipped. Confocal z-stacks were acquired (LSM 700, Zeiss, Germany) and analyzed with ZEN software (Zeiss, Germany).

#### Estimation of the number and strength of fibers

The number and strength of MNTB fibers synaptically connected to the recorded LSO neurons were estimated using a Gaussian mixture model (Fig. 2), k–means clustering (Fig. S2), and density-based spatial clustering of applications with noise (DBSCAN; Fig. S3). The same data set was used in all three approaches.

##### Gaussian mixture model

The number of MNTB inputs was estimated from the stimulus-response plot (Fig. 2B) by fitting Gaussian mixture models (GMM) to the postsynaptic peak amplitudes. GMMs model the observed distribution of PSC peak amplitudes as arising from a linear combination of peak amplitudes of constituent fibers (each following a Gaussian distribution). We used the function `fitgmdist` in MATLAB (MathWorks, R2022a), which uses an expectation maximization (EM) algorithm, to estimate the parameters (mean, variance, and mixing coefficients) of these constituent Gaussian distributions. The algorithm was iterated until the increase of the posterior log-likelihood function value fell below 10^-6^, up to a maximum of 1000 iterations. All fits to our data converged in fewer than the maximum number of iterations. For each model containing a varying number of Gaussian(s) (1–20), GMM fitting was performed 100 times, using the K-means++ algorithm to initiate from random and well-separated Gaussians on each repetition to mitigate the sensitivity of EM to the initiation parameters. The resulting 2000 models were evaluated using the Bayesian information criterion (BIC) score computed using the MATLAB `fitgmdist` function (Schwarz, 1978). A lower BIC score provides stronger support for a given model, but the minimum BIC score has the downside of underestimating the complexity of the data (1). Therefore, among the models with a BIC score up to 20 above the minimum, we considered the one with the most Gaussians as the optimal model, for providing a balance between underestimation and overestimation of the complexity of data (2–4). From the optimal model, the number of MNTB fibers was estimated as the number of Gaussians (the number of steps in peak amplitudes; Fig. 2B), and fiber strength was measured as the distance between the means of neighboring Gaussians. Maximum amplitude was measured from the mean of the five largest peak amplitudes. The coefficient of variation (CV) shown in Fig. 2J was calculated from all fiber strengths for each LSO neuron.

##### K-means clustering

Peak amplitudes of MNTB-evoked postsynaptic responses were subjected to unsupervised, iterative k-means clustering (Fig. S2). K-means clustering was performed for predefined cluster numbers (k = 1-12) for a maximum of 100 iterations with the Euclidean distance metric and k-means++ initialization using the `kmeans` function in MATLAB (R2024b). Clustering quality was assessed using the elbow method with the within-cluster sum of squares (WCSS) and the mean silhouette score computed with MATLAB’s `silhouette` function. The elbow method with the WCSS was used to evaluate the possibility of a single cluster (k = 1). A single cluster was considered optimal when the reduction in WCSS from k = 1 to k = 2 was no more than 50%, indicating no meaningful partitioning. With our data set, this was never the case, and the optimal cluster model (k > 1) was defined as the model yielding the highest mean silhouette score, reflecting maximal within-cluster cohesion and between-cluster separation. From the optimal cluster model, the number of input fibers was determined as the number of clusters (k), and the fiber strength was computed as the distance between the centroid of neighboring clusters.

##### Density-based spatial clustering of applications with noise (DBSCAN)

Peak amplitudes of MNTB-evoked postsynaptic responses were clustered using density-based spatial clustering of applications with noise (DBSCAN) implemented with the `clusterDBSCAN` function in MATLAB (R2024b; Radar Toolbox, version 24.2). DBSCAN was performed with a predefined minimum number of data points and a systematically determined maximum distance of data points within a cluster (epsilon, ε). The minimum number of points in a cluster was set to 3 to capture a small number of observations of synaptic inputs arising from an individual fiber. The ε was determined based on a sorted distance to the k-nearest neighbor for each data point (k-distance graph), using the `clusterDBSCAN.estimateEpsilon` function in MATLAB (R2024b). Because ε is a critical parameter for defining cluster boundaries, an unbiased procedure was used to screen and determine an optimal ε for each neuron. For the number of nearest neighbors (k) from 2 to 11, ε was calculated as the ε value of the “knee” of each k-distance graph, defined as the point with the largest perpendicular distance from the line connecting the first and last points. Because ε varies with k, the three to ten smallest ε values were averaged (i.e., 8 mean ε’s in total), and the averaged ε’s were used as ε candidates for DBSCAN clustering (see Fig. S3C and D). The eight candidate cluster models were evaluated using silhouette analysis. The ε and the corresponding model that produced the highest silhouette score were considered optimal. If multiple ε values tied for the highest silhouette score, the smallest ε among the tied candidates was selected, favoring a model with more clusters. From the optimal cluster model, the number of input fibers was determined as the number of clusters, and the fiber strength was computed as the distance between the means of neighboring clusters.

#### Peak-scaled nonstationary fluctuation analysis (PS-NSFA)

Miniature IPSCs with rise and decay times < 2 SDs of a neuron’s population mean, baseline currents < 4 SD, and inter-intervals >20 ms were included in the analysis. Traces that showed visual signs of two overlapping events (inflection points in the rise period or multiple peaks) were excluded. mIPSC traces were analyzed by means of PS-NSFA similarly to that described previously (5). mIPSC traces were aligned to the steepest rise time using the first derivative of a smoothed waveform (3-point moving average), and then averaged to obtain a mean waveform. The mean waveform was then peak-scaled to each individual mIPSC. The decay phase of the mean waveform was divided into 100 amplitude bins, and for each bin, the difference between the mean current of each individual trace and the mean current of the scaled waveform was calculated. The variance of these differences was calculated across individual events for each neuron (Fig. 3). To decontaminate the variance arising from stochastic channel closing from baseline noise, baseline variance, measured by averaging variances of 5-ms-long epochs in every inter-event period longer than 200 ms, was subtracted from the variance during the decay phase. The variance during the decay phase was plotted against the amplitude of the decay of the mean waveform (Fig. 3I). A parabola was fitted to the first 75% of bins to avoid skewness towards the peak amplitude (`polyfit` function in MATLAB). Single-channel currents were inferred from the parabola fit using the equation:

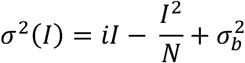

where σ^2^ = variance, *I* is the mean current, *i* is the single-channel current, *N* is the number of open channels at peak current, and σ*_b_*^2^ is background variance. Single-channel conductance (g) was converted from the inferred single-channel current using the reversal potential (Erev, -18.3mV, calculated with the Nernst equation g = I/(Vm – Erev). The number of open receptors was estimated as the median amplitude of the mIPSCs (I) included in the PS-NSFA analysis divided by the single-channel current (i) (Fig. 3J, K).

#### Estimation of the readily releasable pool (RRP)

To estimate the readily releasable pool, we use the train method described by (6). In brief, MNTB fibers were stimulated with a train of 50 stimuli at 100 Hz and for each neuron, the normalized IPSC peak amplitudes were summed in a cumulative normalized IPSC amplitude curve (Fig. 4). A regression line was then fitted to the last 5 points of the cumulative normalized IPSC curve and back-extrapolated to the y-axis. Under the assumption that the replenishment rate is constant throughout the train and that the last responses in the train, after a depletion of the RRP, rely entirely on vesicle replenishment, the y-axis intercept of the linear fit indicates the RRP, and the slope reflects the rate of replenishment. The P_r_ was calculated by dividing the first IPSC amplitude by the RRP. To estimate the number of vesicles in the RRP, the RRP current was divided by the current generated by the release of a single vesicle, estimated from the mean mIPSP amplitude (Fig. 3D). This assumes that the quantal size of a spontaneously released vesicle is the same as that of an action potential-released vesicle, i.e., that vesicles come from the same RRP (7).

**Table S1.**
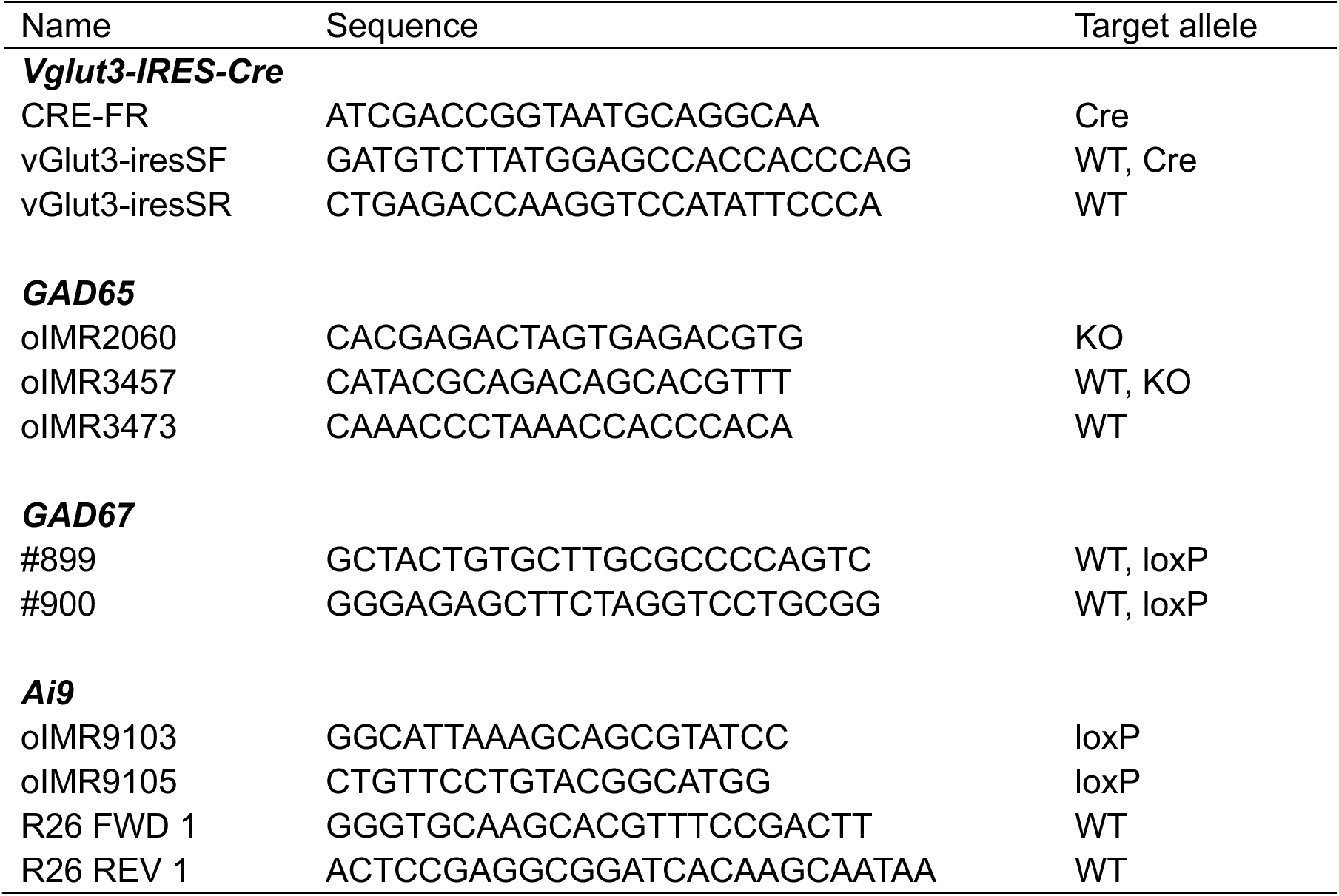
Primer sequences for genotyping.

**Table S2.** Chemical sources.

| Chemical name | Supplier | Catalog number |
| --- | --- | --- |
| CaCl<sub>2</sub> | Sigma-Aldrich | C7902 |
| CNQX | Tocris Bioscience | 1045 |
| CsCl | Acros Organic | 189500500 |
| CsOH | Sigma-Aldrich | C8518 |
| D-gluconic acid | Acros Organic | 119890010 |
| Dextrose | Fisher Scientific | D16 |
| EGTA | Acros Organic | 409910250 |
| HEPES | Sigma-Aldrich | H3375 |
| KCl | Fisher Scientific | P217 |
| KH<sub>2</sub>PO<sub>4</sub> | Fisher Scientific | P5655 |
| Mg-ATP | Sigma-Aldrich | A9187 |
| MgCl<sub>2</sub> | Sigma-Aldrich | M2670 |
| MgSO<sub>4</sub> | Fisher Scientific | M63 |
| Na-GTP | Acros Organic | 226250500 |
| Na<sub>2</sub>-phosphocreatine | Sigma-Aldrich | P7936 |
| NaCl | Fisher Scientific | S271 |
| NaHCO<sub>3</sub> | Fisher Scientific | S233 |
| QX-314 | Tocris Bioscience | 2313 |
| Strychnine | Tocris Bioscience | 2785 |

**Fig. S1.**
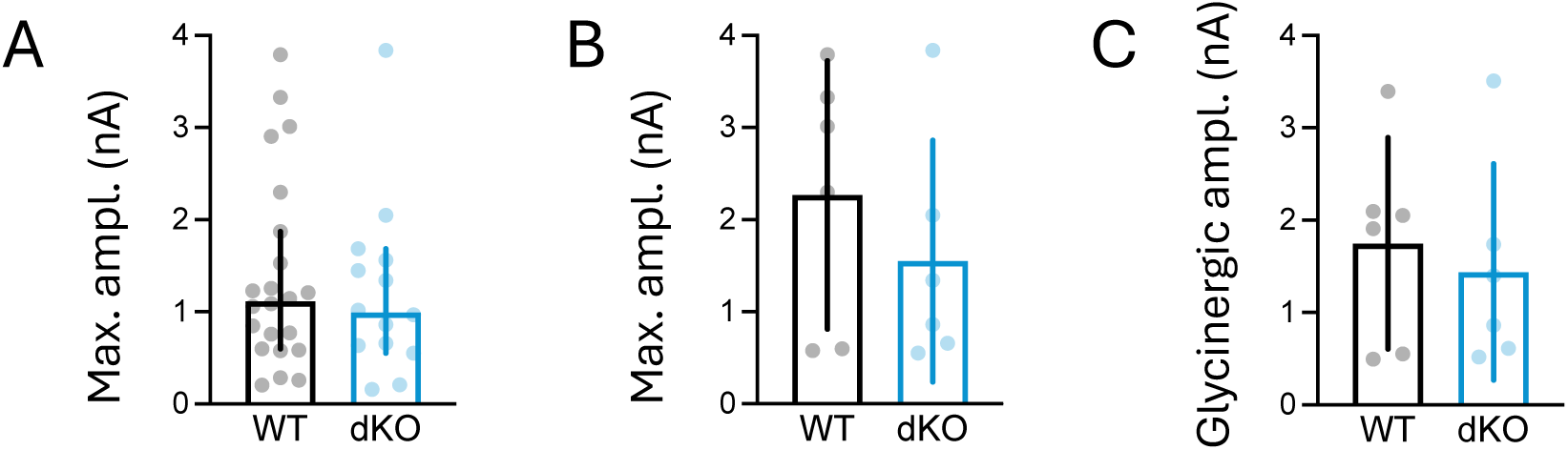
The amplitudes of MNTB-elicited IPSPs did not differ between WT and dKO mice at P3-5. **(A)** Peak amplitudes of MNTB-elicited IPSCs elicited at maximal stimulation without pharmacological blockage (WT: 1.12 (0.60, 1.87) nA (median, 95% CI), n=22, N=14; dKO: 0.99 (0.55, 1.68) nA (median, 95% CI), n=14, N=7; Mann–Whitney *U* = 144.00, *p* = 0.76) **(B)** Same as (A) for neurons shown in Fig 1F. WT: 2.27 ± 1.46 nA (mean, 95% CI), dKO: 1.55 ± 1.31 nA (mean, 95% CI), *t_(10)_* = 0.94, *p* = 0.37, unpaired Student’s t-test) **(C)** No difference between WT and dKO in the pharmacologically isolated glycinergic IPSPs of the neurons shown in Fig 1F (WT: 1.75 ± 1.15 nA (mean, 95% CI) dKO: 1.44 ± 1.17 nA (mean, 95% CI); *t_(10)_* = 0.49, *p* = 0.64, unpaired Student’s t-test)

**Fig. S2.**
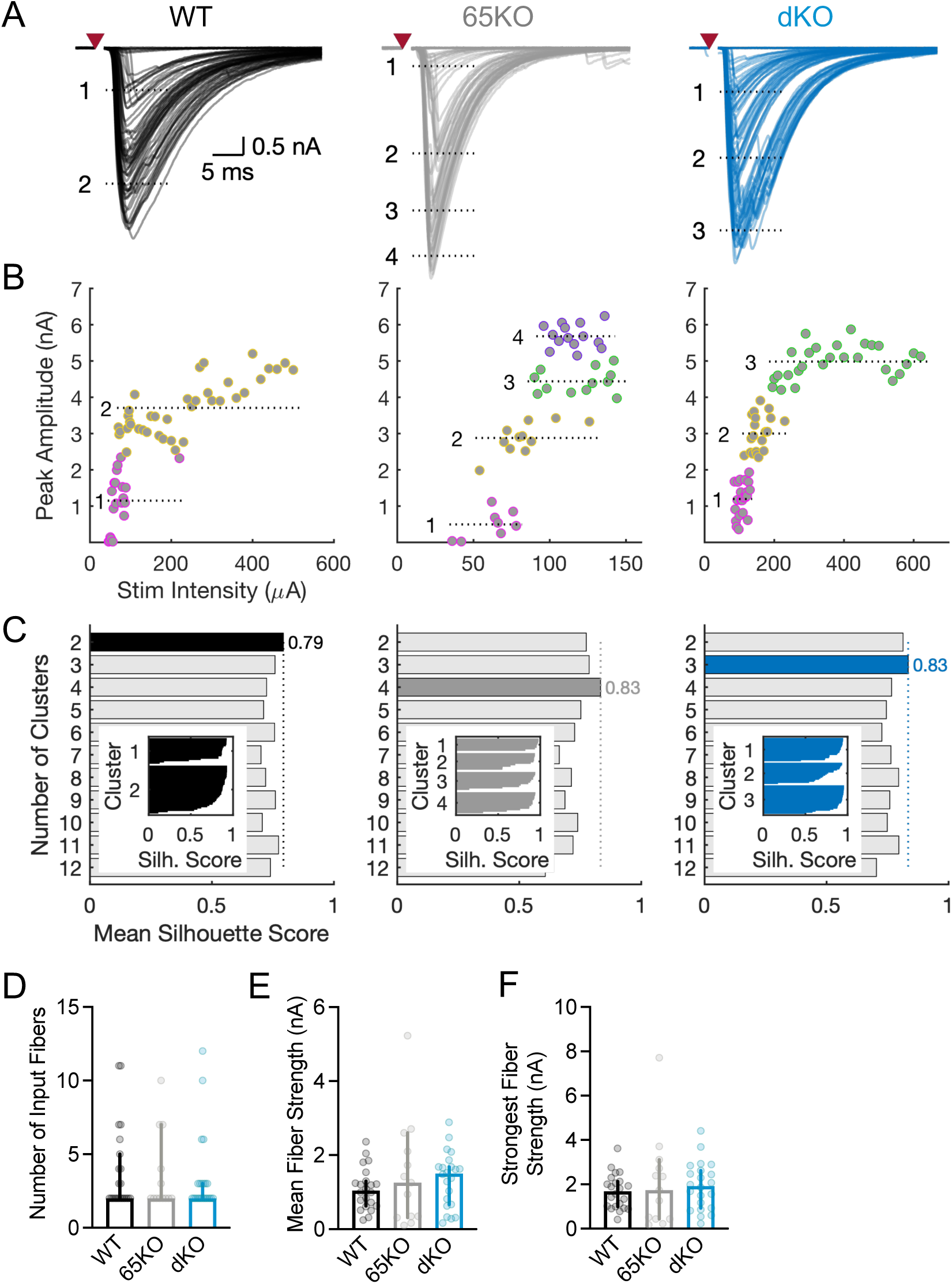
No difference in the developmental pruning and strengthening of MNTB-LSO connections between genotypes indicated by k-means clustering. **(A)** Superimposed traces of the synaptic currents recorded in an example LSO neuron from each genotype (left to right: WT, 65KO, dKO, same neurons as in figure 2) in response to electrical stimulation with gradually increasing stimulation currents. Red triangles indicate time the of stimulation. Stimulus artifacts are omitted for clarity. Dashed lines indicate the mean of each cluster. Clusters correspond to those in (B) and (C) **(B)** Plots of peak amplitude versus stimulation intensity for the three example neurons in (A). Amplitudes are clustered using k-means and indicated by different colors. Dashed lines indicate means of clusters. The distance between the means of neighboring clusters indicates fiber strength. **(C)** Mean silhouette scores of clustered data obtained by specifying different cluster numbers (ranging from 2 to 12) for the example neurons. The cluster number with the highest mean silhouette score was considered the optimal number for each neuron. **(Inset)** Individual silhouette scores for each cluster of the optimal cluster number. **(D)** Number of input fibers, **(E)** Mean fiber strength, and **(F)** Strongest fiber strength in the population of LSO neurons. None of these parameters were different between genotypes (WT: n = 22, N = 15; 65KO: n = 13, N = 4; and dKO: n = 22, N = 10; number of input fibers: Kruskal-Wallis, *H_(2)_* = 0.19, *p* = 0.91; mean fiber strength: Kruskal-Wallis, *H_(2)_* = 1.31, *p* = 0.51; strongest fiber strength: Kruskal-Wallis, *H_(2)_* = 0.55, *p* = 0.76).

**Fig. S3.**
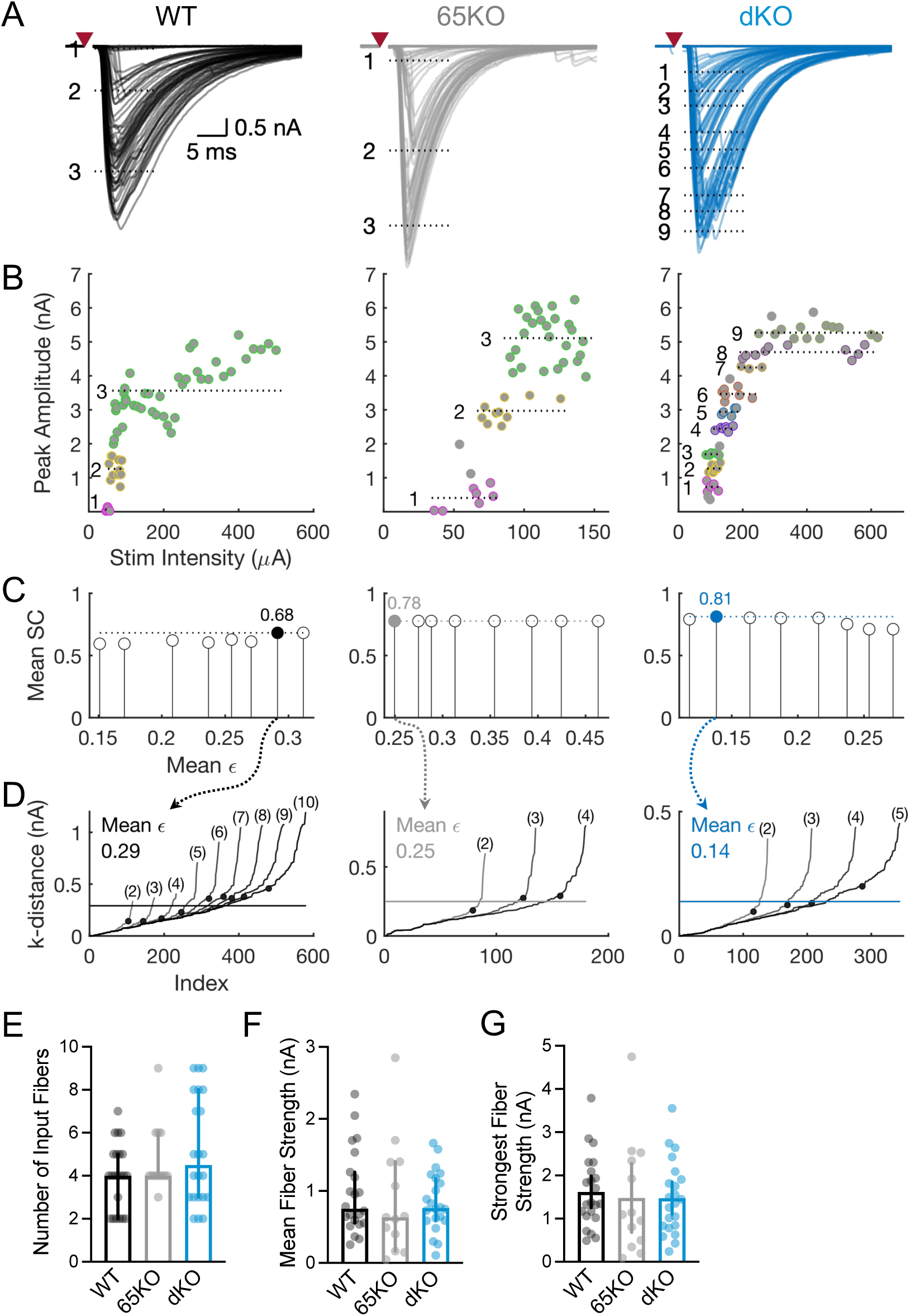
No difference in the developmental pruning and strengthening of MNTB-LSO connections between genotypes indicated by DBSCAN. **(A)** Superimposed traces of the synaptic currents recorded in an example LSO neuron from each genotype (left to right: WT, 65KO, dKO, same neurons as in Fig. 2 and Fig. S1) in response to electrical stimulation with gradually increasing stimulation currents. Red triangles indicate the time of stimulation. Stimulus artifacts are omitted for clarity. Dashed lines indicate the mean of each cluster. Clusters correspond to those in (B). **(B)** Plots of peak amplitude versus stimulation intensity for the three example neurons in (A). Amplitudes are clustered using DBSCAN and indicated by different colors. Grey indicates datapoints not assigned to any cluster (‘noise’). Dashed lines indicate the cluster means. The distance between the means of neighboring clusters indicates the fiber strength. **(C)** Mean silhouette scores (SC) of eight DBSCAN cluster models generated using eight different ε candidates. Each ε candidate value is the mean of three or more smallest ε estimates, which were determined from the k-distance graph for each k-nearest neighbor from 2 to 11 (see panel D and supplemental Methods). The value of ε of the cluster model with the highest mean silhouette score (filled circle, dashed line) was used as the optimal ε for clustering peak amplitudes with DBSCAN. **(D)** k-distance graphs and ε estimates (filled circles) that served to determine the optimal mean ε for the model with the highest silhouette score (filled circle in C). Each k-distance curve represents the sorted distances to the k-nearest neighbors (k parentheses). The ε estimate for each curve was calculated using the ‘knee’ in the elbow method (see supplemental Methods). The number of k-distance curves used to determine the mean optimal ε (horizontal line) varied between neurons, depending on which cluster model had the highest silhouette score. **(E)** Number of input fibers, **(F)** Mean fiber strength, and **(G)** Strongest fiber strength for the population of LSO neurons. None of these measurements were different between genotypes (WT: n = 22, N = 15; 65KO: n = 13, N = 4; and dKO: n = 22, N = 10; number of input fibers: Kruskal-Wallis, *H_(2)_* = 1.88, *p* = 0.39; mean fiber strength: Kruskal-Wallis, *H_(2)_* = 0.65, *p* = 0.72; strongest fiber strength: One-way ANOVA, *F(2,54)* = 0.15, *p* = 0.86.

**Fig. S4.**
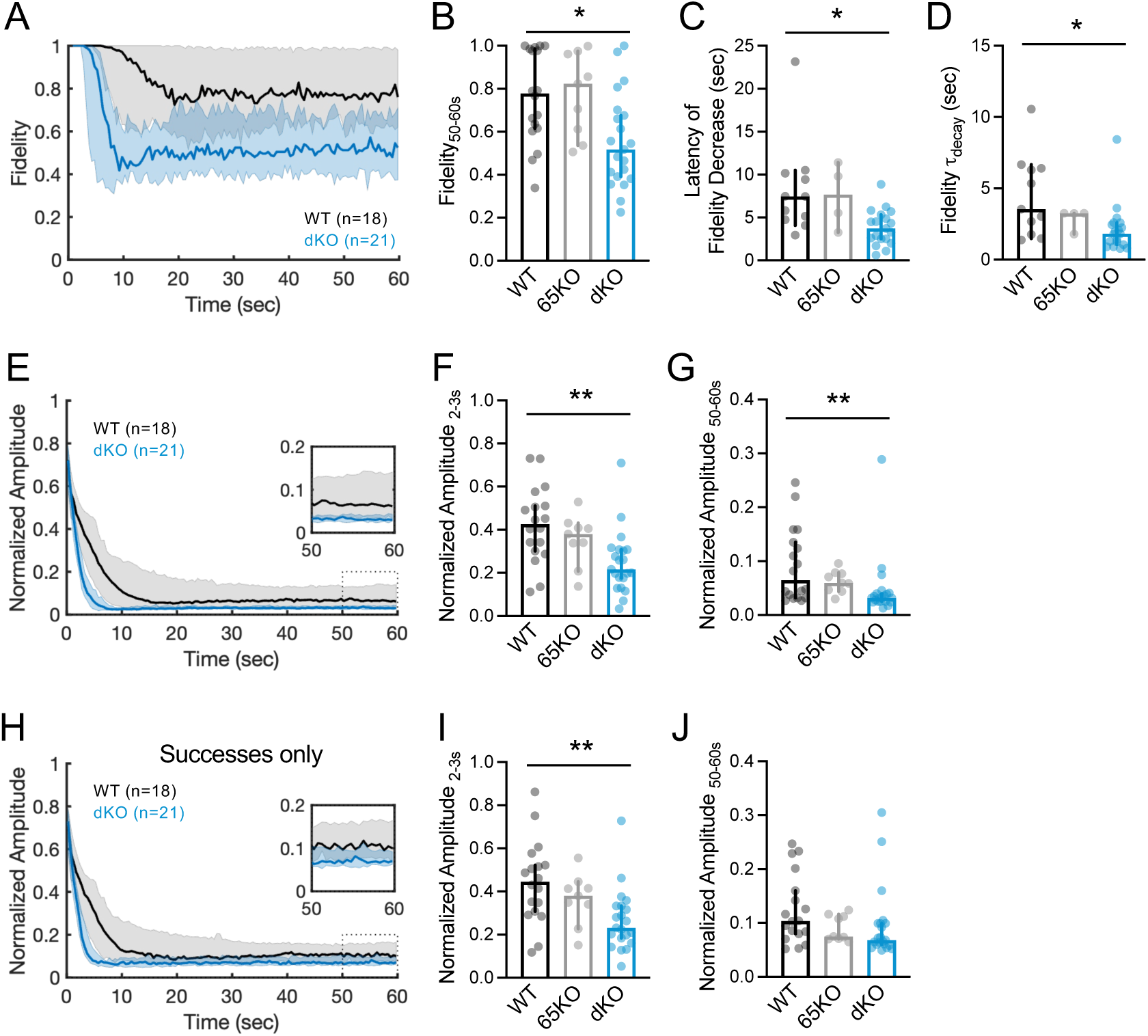
Fidelity and response amplitudes during 50 Hz train stimulation analyzed using an extended response window of 1.5 to 15 ms after stimulus. **(A)** Population plot of fidelity (median ± 95% CI, bin size 0.5 s) for WT and dKO mice. For clarity, data from 65KO mice are omitted. In this and all subsequent panels, the population of neurons is the same as in Fig. 5. **(B)** Fidelity during the last 10 s of the train was significantly lower in dKO mice than in WT mice (Kruskal-Wallis, *H_(2)_* = 9.60, *p* = 0.008; Dunn’s test, WT vs dKO, *p* = 0.014, WT vs 65KO, *p* > 0.99, dKO vs 65KO, *p* = 0.08). (WT: n = 18, N = 8; 65KO: n = 9, N = 6; dKO: n = 21, N = 6). **(C)** Latency to drop of fidelity was significantly shorter in dKO mice than in WT mice (Kruskal-Wallis, *H_(2)_* = 10.07, *p* = 0.007; Dunn’s test, WT vs dKO, *p* = 0.011, WT vs 65KO, *p* > 0.99, dKO vs 65KO, *p* = 0.15). **(D)** Fidelity decrease (tau) was significantly faster in dKO mice than in WT mice (Kruskal-Wallis, *H_(2)_* = 7.95, *p* = 0.019; Dunn’s test, WT vs dKO, *p* = 0.023, WT vs 65KO, *p > 0.99*, dKO vs 65KO, *p* = 0.34). In (C) and (D), neurons with a latency longer than 55 s were excluded (WT, n = 7 out of 18; 65KO, n = 5 out of 9; dKO, n = 2 out of 21). **(E)** Population plot of IPSC amplitudes normalized to the first response (median ± 95% CI bin size, 0.5 s). For clarity, data from 65KO mice are omitted. **(F)** Normalized IPSC amplitudes in dKO mice are significantly reduced 2-3 s after train onset compared to WT mice (Kruskal-Wallis, *H_(2)_* = 11.96, *p* = 0.003; Dunn’s test, WT vs dKO, *p* = 0.002, WT vs 65KO, *p* > 0.99, dKO vs 65KO, *p* = 0.12). **(G)** Steady state depression in dKO mice was stronger than in WT mice (Kruskal-Wallis, *H_(2)_* = 10.73, *p* = 0.005; Dunn’s test, WT vs dKO, *p* = 0.009, WT vs 65KO, *p* > 0.99, dKO vs 65KO, *p* = 0.05). **(H)** Normalized IPSC amplitudes of successful responses only (bin size, 0.5 s). **(I)** Normalized amplitudes in dKO mice are significantly reduced during 2-3 s after train onset compared to WT mice (Kruskal-Wallis, *H_(2)_* = 11.51, *p* = 0.003; Dunn’s test, WT vs dKO, *p* = 0.003, WT vs 65KO, *p* > 0.99, dKO vs 65KO, *p* = 0.16). **(J)** Steady state depression was indistinguishable between genotypes (Kruskal-Wallis, *H_(2)_* = 4.64, *p* = 0.10). \**p* < 0.05, \*\**p* < 0.01.

**Fig. S5.**
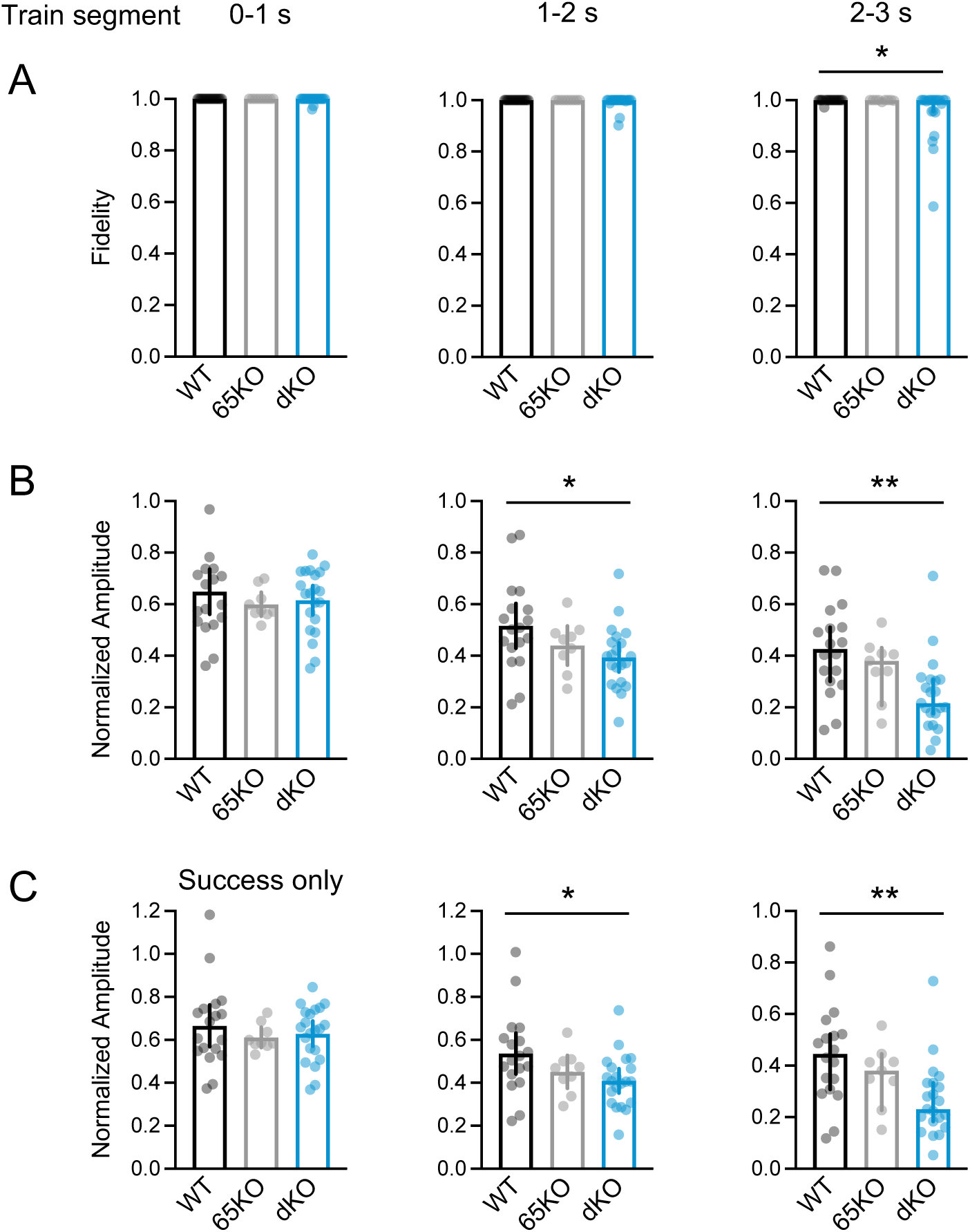
Comparison of fidelity and IPSC amplitudes in the first three seconds of a 50Hz stimulus train. Fidelity **(A)** and the normalized amplitude of all responses **(B)** and successes only **(C)** in the first three seconds of a 50Hz stimulus train. (**B, C**) Each data point represents the average amplitude of 50 responses for each neuron, normalized to the first response in the train. Note the increased synaptic depression of successful responses in dKO mice despite the absence of a significant decrease in fidelity (1-2s). Same cells and analysis as in Fig. S4.

**Fig. S6.**
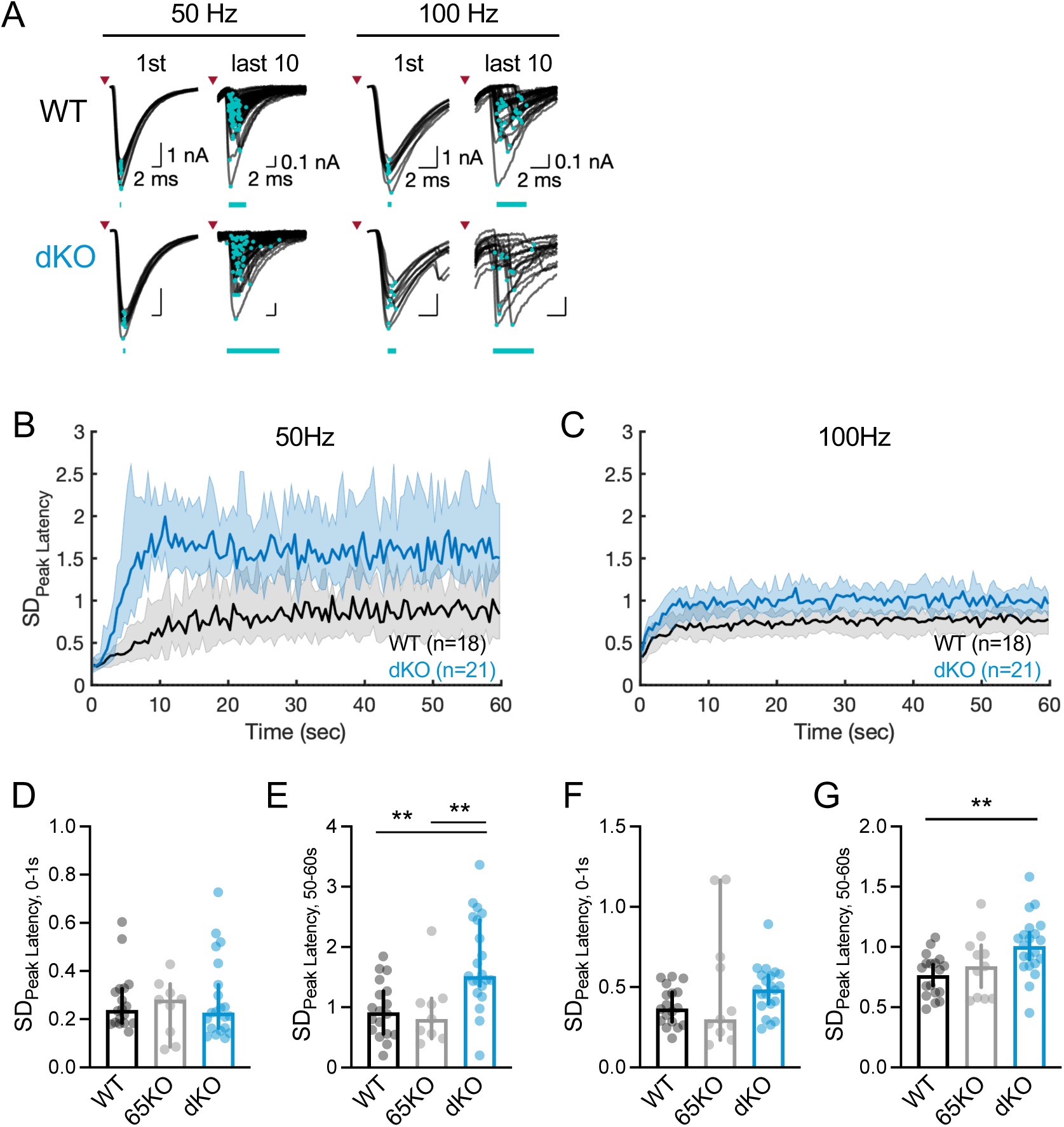
Temporal precision of synaptic transmission during stimulus trains at 50 Hz and 100 Hz, analyzed using an extended response window (time window after stimulus: 1.5 to 15 ms for 50 Hz, and 1.5 to 8 ms for 100 Hz). **(A)** Superimposed traces of successful postsynaptic responses to the first and last ten stimulations at 50-Hz (left) and 100-Hz (right) in an example LSO neuron from a WT mouse (top) and a dKO mouse (bottom). Peaks are marked by turquoise dots, and their jitter is indicated by turquoise bars. Example neurons are the same as in Fig. 6 **B)** Standard deviation of peak latency (median and 95% CI) throughout stimulus trains (0.5 s bins) at 50 Hz, and **(C)** at 100 Hz in WT and dKO mice. To account for fewer successful events in dKO mice (Fig. S4 A,B), SD_Peak Latency_ was computed from 100 bootstrapped successful responses in each 0.5-s bin. For clarity, 65KO data are not shown in (B) and (C). The increase in in SD_Peak Latency_ was significantly larger in dKO mice (50 Hz: Two-way RM ANOVA, *F_(12.7, 569.9)_* = 18.05, *p* < 0.001, time; *F_(2, 45)_* = 11.28, *p* < 0.001, genotype; *F_(25.33, 569.9)_* = 2.12, *p* = 0.001, interaction; WT; n = 18, N = 8, 65KO: n=9, N=6; dKO: n = 21, N = 6); 100 Hz: Two-way RM ANOVA, *F_(20.2, 480.8)_* = 14.64, *p* < 0.001, time; *F_(2, 47)_* = 7.14, *p* = 0.002, genotype; *F_(20.46, 480.8)_* = 1.01, *p* = 0.45, interaction, WT: n = 18, N = 8; 65KO: n = 11, N = 7; dKO: n = 21, N = 6). **(D)** SD_Peak Latency_ in the beginning (0-1 s), and (**E**) at the end (50-60 s) of 50-Hz trains. At steady state, SD_Peak Latency_ of dKO mice was larger than in WT and 65KO mice (Kruskal-Wallis, *H_(2)_* = 14.01, p < 0.001; Dunn’s test, WT vs dKO, p = 0.004, WT vs 65KO, p > 0.99, dKO vs 65KO, p = 0.010). (**F, G**) Same as (D,E) but for 100 Hz trains. At steady state, SD_Peak Latency_ was significantly larger in dKO mice than in WT mice (one-way ANOVA, *F_(2, 47)_* = 5.82, *p* = 0.006; Tukey’s test, WT vs dKO, *p* = 0.005, WT vs 65KO, *p* = 0.67, dKO vs 65KO, *p* = 0.13). **\***\**p* < 0.01.

## Datasets and software

Datasets for supplementary figures and MATLAB scripts are available at https://github.com/jwlee330/GABA_co-release/

